# Detecting rapid pan-cortical voltage dynamics in vivo with a brighter and faster voltage indicator

**DOI:** 10.1101/2022.08.29.505018

**Authors:** Xiaoyu Lu, Yunmiao Wang, Zhuohe Liu, Yueyang Gou, Dieter Jaeger, François St-Pierre

## Abstract

Widefield imaging with genetically encoded voltage indicators (GEVIs) is a promising approach for understanding the role of large cortical networks in the neural coding of behavior. However, the slow kinetics of current GEVIs limit their deployment for single-trial imaging of rapid neuronal voltage dynamics. Here, we developed a high-throughput platform to screen for GEVIs that combine fast kinetics with high brightness, sensitivity, and photostability under widefield one-photon illumination. Rounds of directed evolution produced JEDI-1P, a green-emitting fluorescent indicator whose performance is improved for all metrics. Next, we optimized a neonatal intracerebroventricular delivery method to achieve cost-effective and wide-spread JEDI-1P expression in mice. We also developed an approach to effectively correct optical measurements from hemodynamic and motion artifacts. Finally, we achieved stable brain-wide voltage imaging and successfully tracked gamma-frequency whisker and visual stimulations in awake mice in single trials, opening the door to investigating the role of high-frequency signals in brain computations.

## Introduction

Dissecting the neural underpinnings of behavior has been a long-pursued aim in neuroscience. Because many brain functions require coordinated computations between brain regions, they cannot be understood from studying brain areas in isolation. Therefore, capturing brain-wide activity and interactions is an increasingly common approach in systems neuroscience ^1,2^. For example, monitoring large-scale neural activity can be used to determine the distributed representations of task features and decisions ^3^. Several techniques have been applied for brain-wide recording of neural activity. NeuroPixel probes have been used to monitor activity from thousands of neurons across many brain regions ^4,5^. However, these probes typically cannot distinguish between genetically defined cell populations, produce pan-cortical spatial maps of activity, and report subthreshold membrane voltage fluctuations. Although spikes and local field potentials can be separated with silicon probe recordings, the latter presents a complex mixture of sinks and sources from multiple cell types and layers ^6^.

Genetically encoded calcium indicators are alternative tools that enable the monitoring of neural activity with cell type specificity ^7,8^. Calcium indicators have been deployed for brain-wide cortical imaging ^9,10^. However, the slow indicator response kinetics and the intrinsically slow dynamics of calcium signals limit the ability of calcium indicators to follow rapid neural activity. Moreover, calcium indicators generally do not report subthreshold depolarizations and hyperpolarizations ^11,12^. Calcium indicators are therefore poorly suited to report critical oscillatory properties of neural population activity, such as 15-30 Hz beta-band activity in the control of motor network function ^13–15^ and 40-70 Hz gamma oscillations that may dynamically link functional neural networks in perception and cognition ^16–19^.

Emerging tools for spiking and subthreshold electrical activity measurement are genetically encoded voltage indicators (GEVIs) — sensors that report changes in voltage as variations in fluorescence ^11^. GEVIs have been shown to report spikes, subthreshold depolarizations, and hyperpolarizations in genetically defined cell types and can feature faster temporal precision than calcium indicators ^20–25^. Previous generations of GEVIs have been used for widefield cortical imaging ^12,26–32^ However, while these GEVIs have been used for investigating slow cortical waves in the delta band (0.5-4 Hz) ^29,32,33^ they are suboptimal for following fast voltage changes such as gamma oscillations due to suboptimal for response amplitudes and/or slow kinetics.

Here, we report on a new GEVI optimized for one-photon (1P) imaging and improved methods for brain-wide voltage imaging suitable for chronic recordings in behaving mice. We first discuss the development of an automated high-throughput platform for screening indicators across multiple performance characteristics. This system was utilized to evolve JEDI-1P, a GEVI with improved response amplitude, kinetics, brightness, and photostability. We optimized our genetic strategy to produce robust brain-wide expression of JEDI-1P and developed an optimized approach to correct for hemodynamic and motion artifacts. The improvements in the indicator, expression strategy, and imaging methods enabled long-term brain-wide voltage imaging and successfully tracking high-frequency cortical voltage oscillations in the gamma band for the first time.

## Results

### Enabling multiparametric GEVI screening under widefield one-photon illumination

We first sought to develop a new GEVI based on the ASAP-family of indicators, where a circularly permuted GFP (cpGFP) is inserted in an extracellular loop of a voltage-sensing domain derived from a voltage-sensitive phosphatase ^24^. Voltage-induced conformational changes in the voltage-sensing domain modulate the brightness of the coupled GFP (**Fig. 1a**). We chose to focus on this class of indicators because^34^ they have been shown to express efficiently and report voltage in a variety of neuronal types and experimental preparations ^24,25,35–38^ In addition, cpGFP-based sensors have the potential to produce larger responses than FRET-based indicators ^34^.

**Figure 1.**
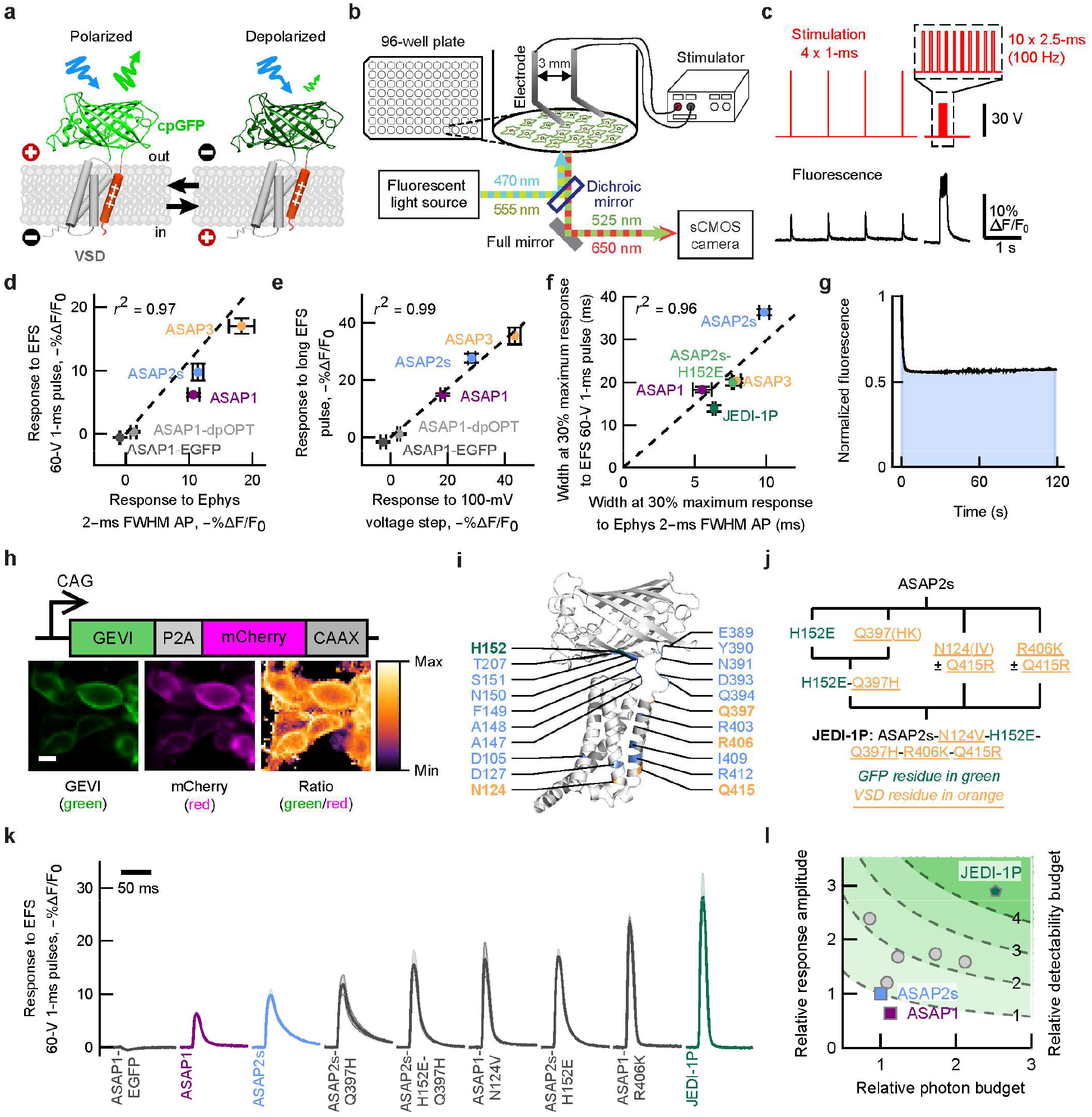
Multiparametric one-photon screening identified JEDI-1P, a voltage indicator with improved performance. a. We engineered GEVIs composed of a circularly permuted green fluorescent protein (cpGFP, green) whose brightness is modulated by voltage-induced conformational changes in the voltage sensitive domain (VSD, gray and orange) b. Schematic of the platform for GEVI screening under widefield one-photon illumination. c. *Top*, Electric field stimulation (EFS) protocol. Inset: zoom-in of the 100-Hz pulse train. *Bottom*, representative fluorescence responses of the parental indicator, ASAP2s. d. Fluorescence responses to 1-ms EFS pulses and AP-like waveforms (2-ms width at half-maximum, +30 mV peak, -70 mV baseline, whole-cell voltage clamp) are highly correlated. For (**d-f**), n = 4 independent transfections (EFS) or 3-11 HEK293A cells (voltage clamp) per GEVI. e. Fluorescence responses to EFS pulse trains and 1-s 100-mV step depolarizations (-70 mV to + 30 mV, whole-cell voltage clamp) are highly correlated. f. Optical response widths to 1-ms EFS pulses and AP-like waveforms are highly correlated. Widths were measured at 30% of the response peak. g. Representative screening data illustrating that photostability was quantified as the as the area under the curve of fluorescence vs. time graphs (shaded area). Fluorescence was normalized to its value at t = 0. The irradiance was ∼33 mW/mm^2^. h. Brightness was evaluated as the GEVI/mCherry (green/red) fluorescence ratio. GEVIs and mCherry were co-expressed using a self-skipping 2A sequence (P2A) to normalize for the expression level (Supplementary Fig. 1c). mCherry was targeted to the plasma membrane via a CAAX motif. i. The 21 locations targeted by mutagenesis are indicated in a predicted 3D structure of ASAP2s. Highlighted residues were mutated in JEDI-1P. j. Key steps of evolutionary path leading to JEDI-1P. Mutations in VSD were screened under both 415R (ASAP1) and 415Q (ASAP2s) contexts. k. JEDI-1P shows faster kinetics and higher response amplitude to 1-ms EFS pulses and kinetics than parental indicators (ASAP1 and ASAP2s) and screening intermediates. l. Illustration of GEVI multiparametric directed evolution. Relative photon budget is defined as the product of the area under the normalized photobleaching curve and the relative brightness. Relative detectability budget is defined as the product of the square root of the relative photon budget and the relative response amplitude. Values relative to that of ASAP2s. Gray circles are screening intermediates. All panels: Error bars or shading denote 95% confidence interval (CI). Pearson’s r^2^ calculated from intercept-free unweighted linear regression of the mean values. More details in Statistical Information.

Indicator properties — response amplitude, kinetics, brightness, and photostability — can be dependent on the illumination modality ^39–41^. Therefore, we sought to develop a platform customized to screen GEVIs for widefield (one-photon) imaging, in contrast to our parallel efforts to screen under two-photon excitation ^42^. Many indicator screening efforts illustrate that mutations can improve a performance metric that was evaluated during screening while impairing others that were not quantified ^21^. To achieve global or ‘holistic’ consideration of a GEVI’s performance, we chose to design a platform that would screen for all key indicator properties in the same assay — response amplitude, kinetics, brightness, and photostability.

The first performance metric we considered was the ability for GEVIs to follow rapid voltage transients such as action potentials (APs) and post-synaptic potentials. We fitted a widefield inverted fluorescence microscope with a camera capable of capturing fluorescence with a temporal resolution of 1 ms per image in a field-of-view of 2048×200 pixels (**Fig. 1b**). We screened indicators using a HEK293 cell line that has a resting membrane potential of ∼–77 mV, similar to that of mammalian cortical neurons ^35,43,44^. 100-ms electric field stimulation (EFS) pulses were previously reported to depolarize these cells ^21,45^ (**Supplementary Fig. 1a**). To benchmark GEVIs’ ability to report faster voltage signals, we shortened the pulse duration to 1 ms (**Fig. 1c**). The resulting GEVIs’ responses had widths down to ∼10 ms (**Supplementary Fig. 1b**). Response amplitudes to AP waveforms and 1-ms EFS pulses were strongly correlated (**Fig. 1d**), suggesting that our platform enables screening for larger responses to fast voltage transients.

We also sought to identify candidates with increased response amplitude to longer voltage signals and variants with faster kinetics, with the intention of combining their respective mutations. We observed that the peak response to a 100-Hz train of ten 2.5-ms pulses was highly correlated with the peak response to 1-s 100-mV depolarization voltage step (**Fig. 1e**) The width of AP waveforms and EFS-induced spikes were also highly correlated (**Fig. 1f**). These results demonstrate we can rank indicators based on their response amplitude to longer voltage signals and their kinetics.

Finally, we evaluated indicators for photostability (**Fig. 1g**) and brightness (**Fig. 1h**). To normalize brightness measurements for well-to-well variations in expression level, we co-expressed the red fluorescent protein mCherry using a self-skipping 2A sequence ^46^ (**Fig. 1h**). mCherry was anchored to the plasma membrane via a CAAX membrane targeting motif ^47^. As expected, the GEVI/mCherry ratio showed less variation from well to well compared with GEVI (green) intensity values alone (**Supplementary Fig. 1c**). Finally, to maximize screening throughput, we excluded wells that did not meet pre-defined quality control standards from further data collection (**Supplementary Fig. 1d**).

### Multiparametric screening identified JEDI-1P, an indicator with improved kinetics, response amplitudes, and photon budget under one-photon illumination

We applied our multiparametric screening platform to screen single-site saturation mutagenesis libraries in 96-well plates. ASAP1 ^24^ and ASAP2s ^35^ as starting templates, since ASAP3 ^25^ had not been reported when this project started. We targeted residues around the chromophore of cpGFP, in conserved positions of the voltage-sensing domain, and near the junctions between cpGFP and the voltage-sensing domain. We screened over 1,200 mutants from 21 single-site libraries (**Fig. 1i**), and promising mutations were combined (**Fig. 1j-l**). Our best variant includes 5 mutations from ASAP2s. We call this sensor Jellyfish-derived Electricity-reporting Designer Indicator for 1-Photon, or JEDI-1P.

We evaluated the response of JEDI-1P to voltage changes in single human (HEK293A) cells using one-photon widefield imaging and whole-cell voltage clamp. JEDI-1P produced steep responses between -80 and 0 mV, demonstrating it is well adapted to reporting voltage signals within the physiological range of cortical neurons (**Fig. 2a-b**). JEDI-1P produced larger responses to a 100-mV depolarization step from −70 to +30 mV compared with ASAP2s and ASAP3 (**Fig. 2c**). We did not compare JEDI-1P to JEDI-2P, a GEVI evolved for two-photon microscopy, because JEDI-2P has comparable performance to ASAP3 under one-photon widefield imaging ^42^.

**Figure 2.**
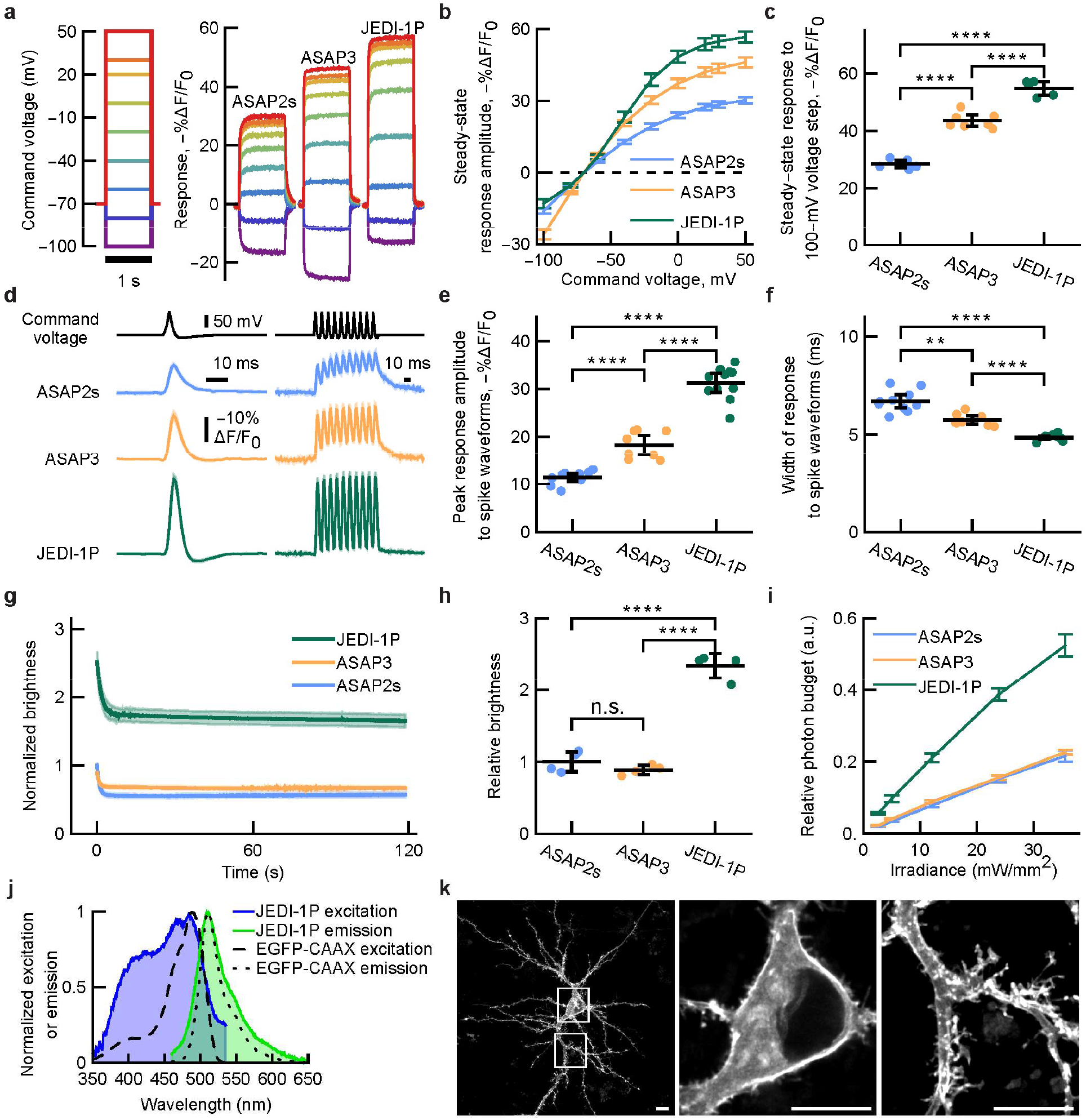
JEDI-1P has improved sensitivity, kinetics, brightness, and photostability under one-photon illumination *in vitro*. a-c. JEDI-1P has greater steady-state response amplitude to 1-s step depolarizations than ASAP2s and ASAP3. Fluorescence was acquired at ∼1 kHz. n = 6 (ASAP2s), n = 7 (ASAP3) and n = 5 (JEDI-1P) HEK293A cells. More details in Statistical Information. a. Mean response to voltage steps. b. F-V curve. c. Responses to 100-mV step voltages (−70 mV to +30 mV). p < 0.0001 (ANOVA). d-f. JEDI-1P shows larger and faster responses to short voltage transients compared with ASAP2s and ASAP3. Fluorescence was acquired at ∼1 kHz. n = 10 (ASAP2s), n = 8 (ASAP3), and n = 11 (JEDI-1P) HEK293A cells. d. *Left*: mean response to spike waveforms simulating APs (2-ms full-width at half-maximum, +30 mV peak, baseline at −70 mV). *Right*: mean response to a 100-Hz spike train waveform. e. JEDI-1P has a larger peak response amplitude to single AP waveforms than ASAP2s and ASAP3. p < 0.0001 (ANOVA). f. JEDI-1P has faster fluorescence responses to single AP waveforms than ASAP2s and ASAP3, as shown by narrower optical responses. The width corresponds to the Full Width at Half Maximum. p < 0.0001 (Welch’s ANOVA), Dunnett’s T3 multiple comparisons test. g. Fluorescence trace of JEDI-1P, ASAP2s, and ASAP3 illuminated with 470/24-nm light at an irradiance of 2.6 mW/mm^2^ at the sample plane, normalized to the fluorescence of the first frame. h. JEDI-1P is brighter than ASAP2s and ASAP3. Brightness was quantified as described in Fig. 1h. n = 4 independent transfections per GEVI. p < 0.0001 (ANOVA). i. JEDI-1P displays larger relative photon budget (area under the curve of the normalized brightness timecourses from panel g) than ASAP2s and ASAP3 over a >10-fold range of irradiance levels (2.6–35.6 mW /mm^2^). j. JEDI-1P has an excitation peak at 484 nm and an emission peak at 509 nm. n = 11 (EGFP-CAAX) and n = 12 (JEDI-1P) independent transfections in HEK293-Kir2.1 cells. k. JEDI-1P efficiently traffics to the plasma membrane in the soma and dendrites. Representative confocal image acquired from a DIV14 rat cortical neuron. All panels: **** p < 0.0001; *** p < 0.001; ** p < 0.01; * p < 0.05; n.s. p > 0.05. Unless otherwise noted, Tukey’s HSD multiple comparison test was used. Error bars or shading denote 95% confidence interval (CI) of the mean.

At ∼33°C, JEDI-1P had on- and off- kinetics of 0.54 ± 0.07 and 1.20 ± 0.12 ms (mean ± SEM), respectively, faster than ASAP2s and ASAP3 (**Table 1**). As predicted from its larger response amplitudes and faster kinetics, JEDI-1P exhibited responses to spike waveforms 174% [148-202%] and 71% [50-94%] (mean, [95% CI]) larger than ASAP2 and ASAP3, respectively (**Fig. 2d-e**). Because of its faster kinetics, JEDI-1P fluorescence recovered to baseline between individual spikes of a train of action potential waveforms (**Fig. 2d**) and produced narrower responses to spike waveforms (**Fig. 2f**).

**Table 1.**
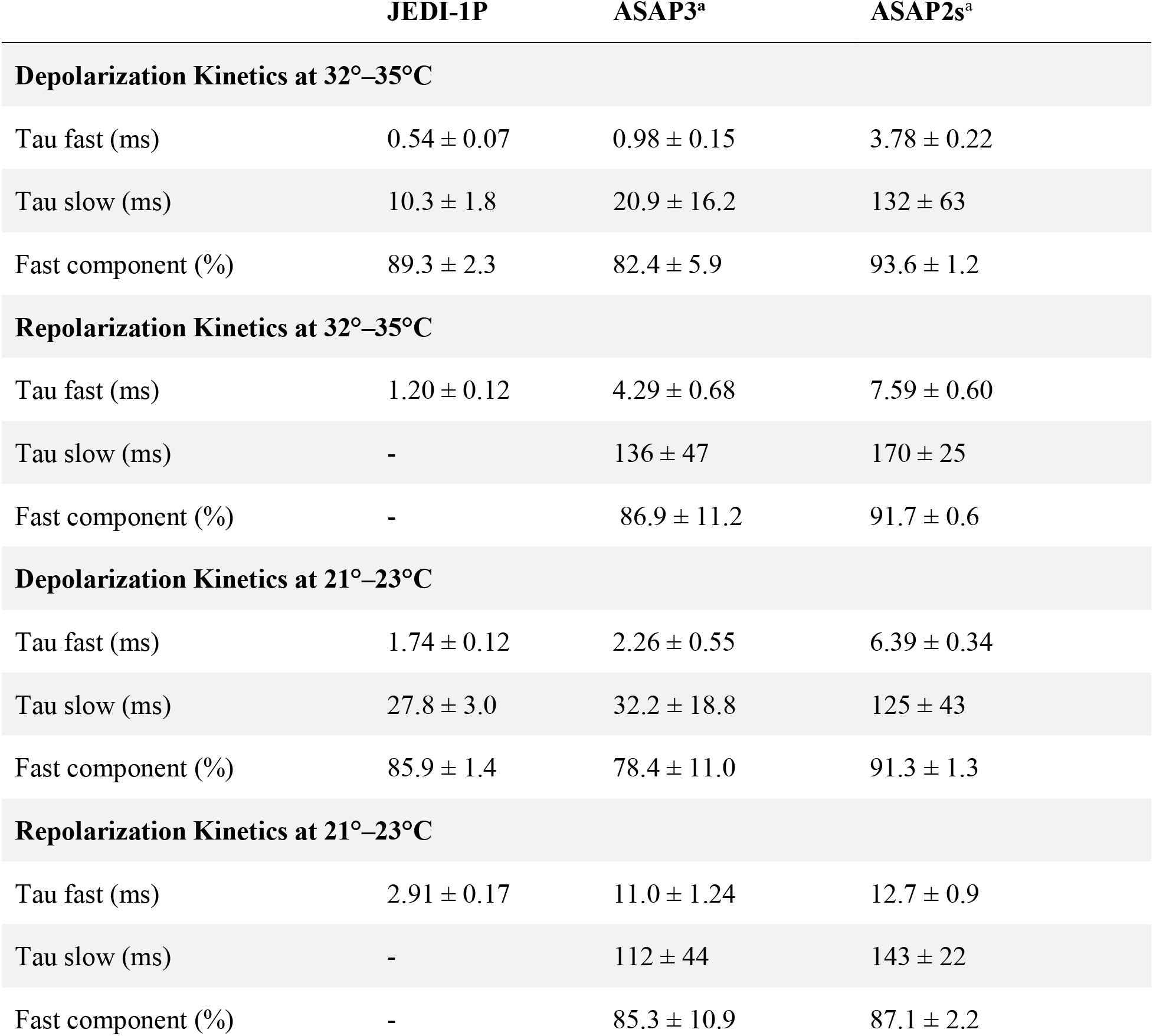
JEDI-1P is faster than ASAP2s and ASAP3, related to Figure 2. Kinetics were measured in response to 1-s voltage steps in HEK293A cells under whole-cell voltage clamp. Cells were clamped at −70 mV at rest. Depolarization was performed from −70 mV to 30 mV while repolarization was performed from 30 mV to −70 mV. Values of JEDI-1P are means ± 95% CI from n = 9 cells at 21-23°C and n = 10 cells at 32-35°C. ^a^ Values from ^80^. Experiments were performed under the same condition as JEDI-1P. Values are means ± 95% CI from n = 9 cells (ASAP2s), n = 9 cells (ASAP3) at 21-23 °C and n = 10 cells (ASAP2s), n = 16 cells (ASAP3) at 32-35 °C.

We next sought to evaluate JEDI-1P’s brightness and photostability. Under illumination at ∼3 mW/mm^2^, indicators exhibited high photostability after an initial ∼5-s period of fast photobleaching (**Fig. 2g**). Compared with ASAP3 and ASAP2s, JEDI-1P had higher brightness at the beginning (**Fig. 2h**) and throughout (**Fig. 2g**) the experiment. As a result of its high photostability and greater brightness, JEDI-1P exhibited a larger photon budget (area under the curve of the relative brightness trace) at all tested irradiance levels (**Fig. 2i**).

To help determine optimal voltage imaging parameters, we acquired JEDI-1P’s excitation and emission spectra. JEDI-1P was maximally excited at 484 nm and produced an emission spectrum with a peak at 509 nm, similar to EGFP (**Fig. 2j**). JEDI-1P can thus be imaged under similar optical configurations as popular GFP-based sensors such as GCaMP ^7^. Finally, we confirmed that JEDI-1P could be expressed and efficiently trafficked to the plasma membrane in the soma and dendrites of cortical neurons *in vitro* (**Fig. 2k**), similarly to its parental sensor ASAP2s ^35^.

### Robust pan-cortical expression of soma-targeted JEDI-1P

To conduct pan-cortical voltage imaging, we first set to optimize JEDI-1P expression across the cortex. To preferentially image somatic voltage signals, JEDI-1P was appended with a motif from the potassium channel Kv2.1 that restricts membrane protein expression to the soma and proximal dendrites ^48^. The resulting variant, JEDI-1P-Kv, was cloned in a Cre-dependent expression vector. JEDI-1P-Kv adeno-associated viruses (AAVs) were packaged using PHP.eB, a capsid that produces efficient gene transfer across the central nervous system ^49,50^. AAVs were injected in *EMX1-cre* mice to obtain preferential JEDI-1P-Kv expression in cortical excitatory neurons ^51,52^.

Total hemoglobin fluctuations and differential light absorption of oxygenated and deoxygenated hemoglobin produce artifactual signals during widefield imaging of neural activity with optical indicators ^53–59^. Imaging a reference fluorescence signal at a different emission wavelength is a common and effective approach to isolate neural signals from hemodynamic activity ^56,58–60^. Therefore, to correct JEDI-1P-Kv measurements from hemodynamic and motion artifacts, we co-injected AAVs expressing the red fluorescent protein (RFP) mCherry from the pan-neuronal promoter *hSyn* (or tdTomato from the non-cell-type-specific promoter *CAG*).

We chose to perform neonatal intracerebroventricular (ICV) ^**61**^ over retro-orbital ^**49**^ injections given the 40-fold higher cost of the latter method. For imaging, we used a clear skull preparation that leaves the skull intact ^**62**,**63**^. Four weeks after neonatal ICV injections, we observed strong JEDI-1P-Kv expression throughout the cortex and in *EMX1*-expressing subcortical structures such as the hippocampus (**Fig. 3a**). As expected, JEDI-1P-Kv expression was limited to somatic and proximal dendritic membranes. mCherry or td-Tomato were also expressed cortex-wide, but in the cytosol of a distinct subset of neurons (**Fig. 3a**, right for mCherry, **Supplementary Fig. 2g** for tdTomato). We imaged the intact cortex of head-fixed mice at 200 Hz (**Fig. 3b**). While there were intraindividual and interindividual differences in fluorescence across the cortex and between individuals, JEDI-1P-Kv could be reliably detected above background fluorescence (**Fig. 3c-d**). Given the strong fluorescence of mCherry, reliable signals were obtained with the same 466/40-nm-filtered blue excitation light source as used for JEDI-1P-Kv (**Fig. 3e**). Using a single light source obviates the need for temporal switching between channels and eliminates differential light noise.

**Fig 3.**
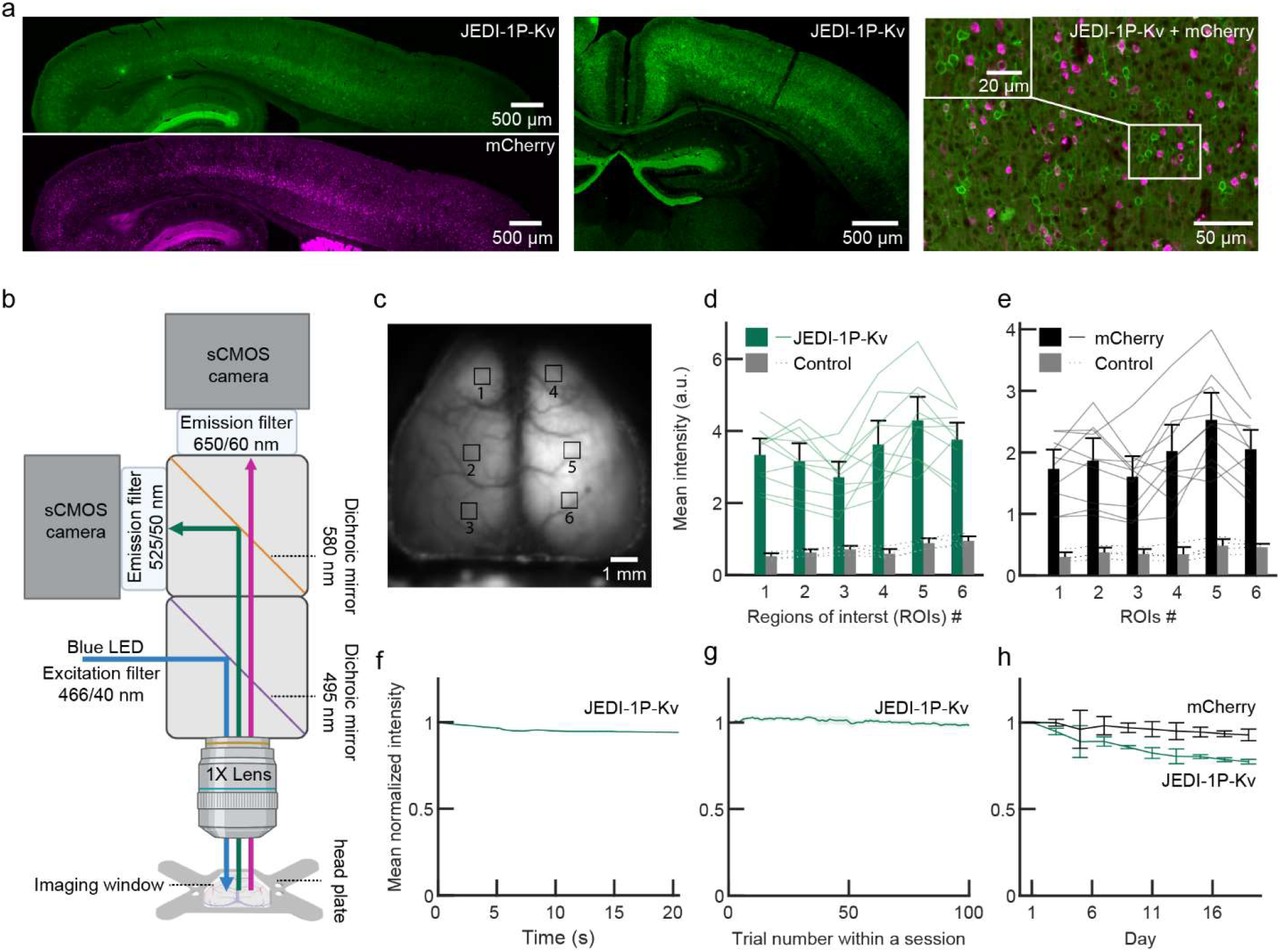
Reliable pan-cortical expression of JEDI-1P-Kv. a. Sagittal slide illustrating that JEDI-1P-Kv (*top left*) and mCherry (*bottom left*) are expressed pan-cortically. *Middle*, coronal section. R*ight*, zoomed-in and overlaid image of JEDI-1P-Kv and mCherry. Adeno-associated viruses encoding soma-localized JEDI-1P (JEDI-1P-Kv) and a cytosolic red-emitting reference fluorescent protein (mCherry) were introduced bilaterally by neonatal intracerebral ventricular (ICV) injection. b. Imaging setup schematic. All experiments were conducted at 200 Hz. c. Representative image of JEDI-1P-Kv acquired with the system depicted in (b). The numbered squares show the 6×6-pixel (600 × 600 μm) regions of interest (ROIs) analyzed in (d) and (e). d-e. JEDI-1P-Kv (d) and mCherry (e) can be robustly detected over the background. Bars represent the mean fluorescence intensity at different ROIs from mice expressing JEDI-1P-Kv and mCherry (n = 10) and control mice expressing neither (n = 4). Lines show data from individual mice expressing JEDI-1P-Kv or mCherry (solid) or control mice (dashed). Mean intensities of all ROIs from JEDI-1P-Kv and mCherry mice are significantly higher than those from control mice. p = 0.002 for each ROI, Mann-Whitney U test. f-h. JEDI-1P-Kv is photostable under wide-field illumination *in vivo*. Excitation power was 0.05 mW/mm^2^, which is lower than used *in vitro* (Fig. 2g-h). Data shows mean JEDI-1P-Kv fluorescence intensity from pixels of cortex over a single trial (f), a daily session (g), and all 10 imaging sessions (h). Between-trial intervals were ∼20 s. Total imaging time per session was 34 min. Data was normalized to the beginning of the trial (f), the mean intensity of the first trial (g), and the mean intensity of the first session of the first day (h). n = 300 trials from 3 mice (100 trials/animal) (f), 27 sessions from 3 mice (g), and 3 mice (h). All error bars and shaded areas denote the 95% CI.

### JEDI-1P-Kv is photostable during widefield longitudinal imaging experiments

GEVI photobleaching can severely limit the duration of voltage imaging sessions with some indicators and experimental preparations ^11,21^. However, we observed minimal photobleaching with JEDI-1P-Kv while conducting widefield imaging (**Fig. 3f-h**). Although the raw fluorescence intensity decayed to 94.1 ± 0.3% (represented as mean ± 95% CI here and henceforth) by the end of a 20-s trial **(Fig. 3f)**, the fluorescence level recovered almost fully during the 20-s dark intervals between trials with 98.4 ± 1.3% intensity remaining at the end of an imaging session (**Fig. 3g**).

After more than 5 h of imaging spread over 19 days — 10 imaging sessions of one hundred 20-s trials — JEDI-1P-Kv conserved 77.4 ± 1.3% of its fluorescence (**Fig. 3h**). Consistent with our 466/40-nm light source exciting mCherry far from its absorption peak of 587 nm ^64^, mCherry displayed even higher photostability than JEDI-1P-Kv by retaining 92.8 ± 3.3 % of its initial fluorescence after the last imaging session. Overall, the high photostability of JEDI-1P-Kv when imaged with our widefield system enabled repeated and longitudinal in vivo optical recordings with high signal-to-noise ratio.

### Data pre-processing pipeline isolates JEDI-1P-Kv voltage signals from hemodynamic artifacts

We developed a pre-processing pipeline that corrects the voltage signals for autofluorescence and photobleaching, and then removes hemodynamic and motion artifacts. Autofluorescence was subtracted using values measured in a group of control mice without AAV injections, and single-trial time courses were corrected for photobleaching using a bi-exponential fit. The remaining signal has a clear heartbeat artifact (**Fig. 4a**), which is also present in the RFP channel. The heartbeat artifact was successfully removed from the voltage signal by regressing the RFP channel filtered around the heartbeat frequency (**Fig. 4a-b**). We achieved best overall correction for signal artifacts by regressing three distinct frequency bands since the voltage signal contamination has different amplitudes at distinct frequencies (**Fig 4c-d, Supplementary Fig. 2a-f**). The number of regression steps and their respective bandwidth can be adjusted depending on the characteristics of motion and hemodynamic contamination in different preparations such as anesthetized vs awake conditions. We also tested the pipeline on data with tdTomato rather than mCherry as the reference protein and achieved similar performance (**Supplementary Fig. 2g-i**).

**Fig 4.**
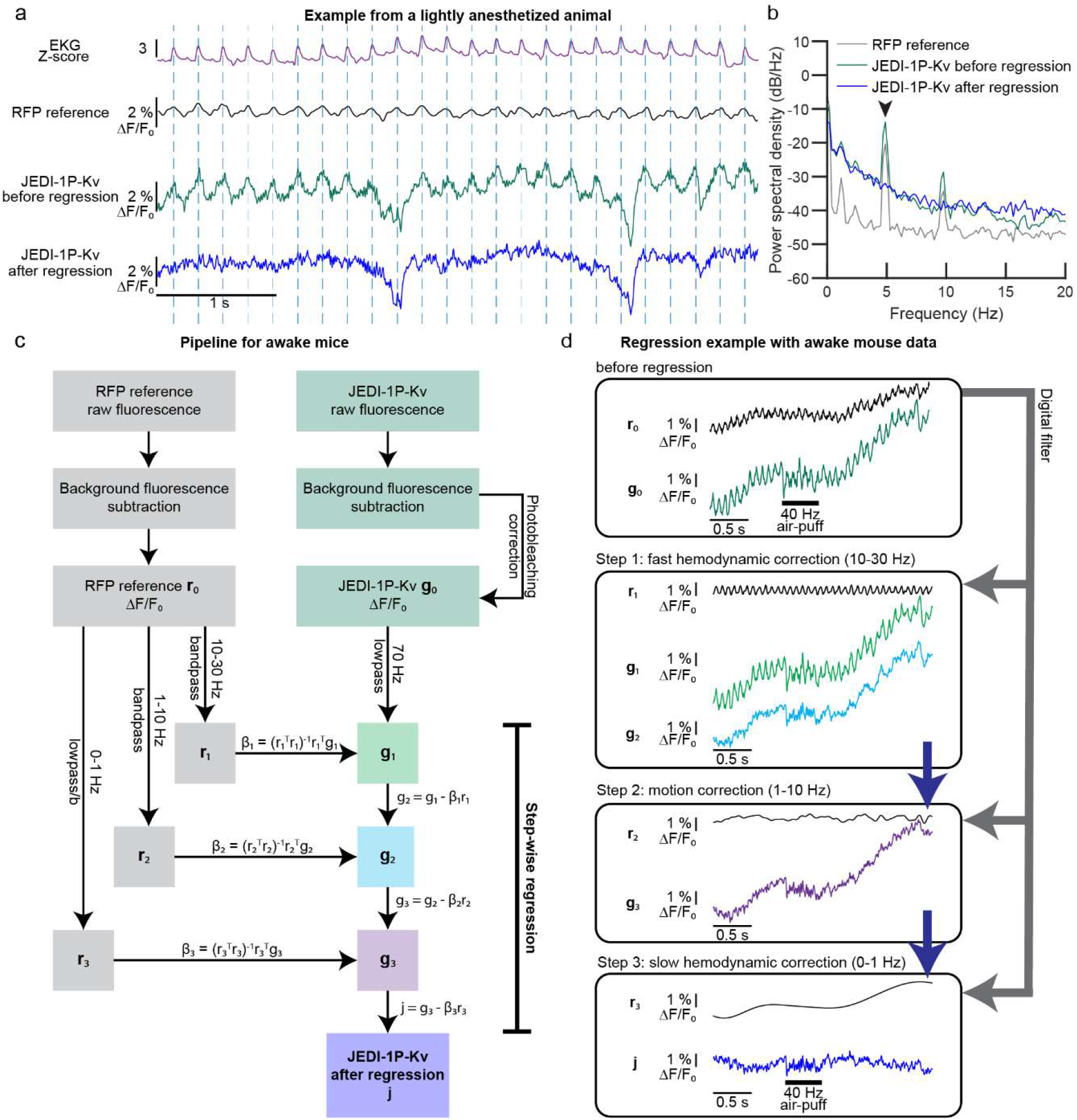
Hemodynamic and motion correction can be achieved with a red fluorescent protein and a single excitation light source. a. 5 Hz oscillations observed in JEDI-1P-Kv and RFP reference channels in lightly anesthetized mice correspond to the heart rate (shown in EKG) and reflect fast blood volume changes (fast hemodynamics). JEDI-1P-Kv and RFP optical traces are from the same 2×2-pixel ROI. The regression pipeline for anesthetized mice is as in panel (c) but used 4-20, 1-4, 0-1 Hz as the three bandpass filters. b. Successful removal of fast hemodynamic signals in a lightly anesthetized animal. n = 25 trials from a representative mouse. The heartbeat peak at ∼5 Hz (arrow) was removed in the regressed trace (blue). c. Signal pre-processing workflow when recording from awake animals. The reference channel was filtered to three different frequency ranges for regression: 10-30 Hz — which includes the heartbeat frequency in the awake condition, 1-10 Hz, and 0-1 Hz. These ranges produced efficient regression in our awake mice, but different ranges might be optimal in other studies. d. Stepwise regression on representative traces removed fluorescence signals unrelated to voltage.

### The relation between JEDI-1P-Kv voltage signal and simultaneous local field potential recordings reveals significant correlations up to 60 Hz

To examine the relation between JEDI-1P-Kv voltage signal and LFP recordings, we performed simultaneous widefield imaging and acute barrel cortex local field potentials (LFPs) recordings in lightly isoflurane-anesthetized mice (**Fig. 5a**). Single-trial optical recordings closely tracked LFPs recorded close to the imaging site (**Fig. 5b**). We quantified the correlation between barrel cortex LFP and JEDI-1P-Kv voltage over the entire cortical area and found that LFP signals were unsurprisingly best correlated with optical responses close to the LFP recording site (**Fig. 5c-d**). In addition, we observed a network spanning bilateral barrel and motor cortices showing significant voltage correlations with barrel cortex LFPs (**Fig. 5c**). This result is consistent with previous fMRI and widefield Ca^2+^ imaging studies showing that LFPs have shared components across functional networks ^56,65,66^. We did not expect correlation coefficients close to the maximum of 1.0 since LFP recordings were not limited to excitatory neurons, correspond to neurons located in different locations and depth, and represent a complex mixture of local potentials and volume conductance from distant sources ^67^.

**Fig 5.**
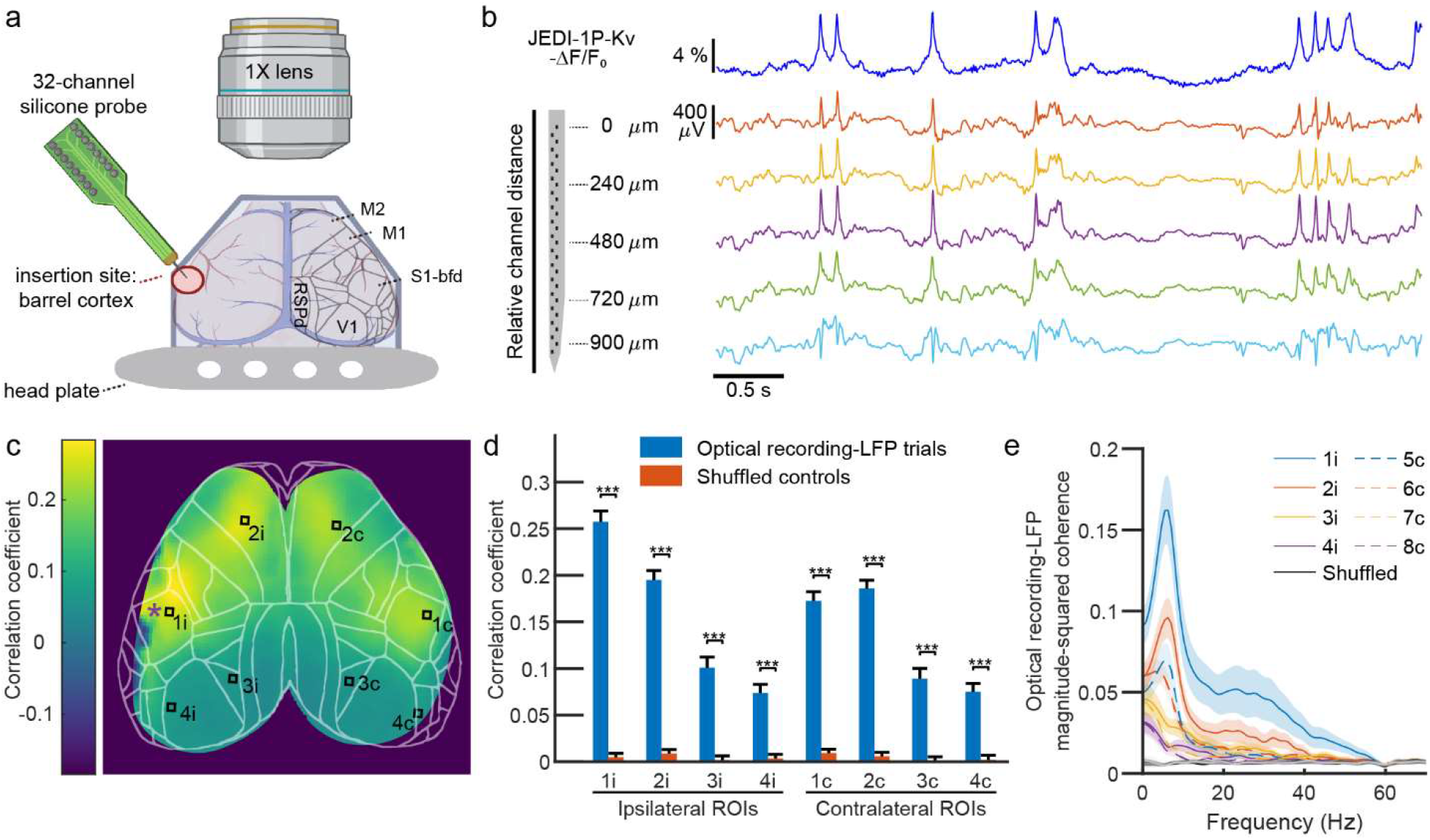
JEDI-1P-Kv signals correlate with local field potentials (LFPs) a. Setup schematic for simultaneous voltage imaging and acute LFP recording setup in lightly anesthetized animals. Lines indicate cortical map derived from the Allen Mouse Brain Common Coordinate Framework ^78^. M1, primary motor cortex. M2, secondary motor cortex; S1-bfd, primary somatosensory (barrel) cortex; RSPd, retrosplenial cortex. b. JEDI-1P-Kv reports voltage transients also seen in LFP traces. Voltage imaging was conducted close to the LFP recording site (ROI 1i in (c)). LFPs were recorded from 32 channels, with traces from 4 channels shown here (channels 1, 9, 17, 25, and 31). Depths are relative to the top channel. c-d. LFPs are correlated with JEDI-1P-Kv signals from both ipsilateral (i) and contralateral (c) sides e. Map showing the mean correlation between the LFP recording and JEDI-1P-Kv traces from 2×2 pixel ROIs in a representative mouse. The purple asterisk indicates the LFP electrode insertion site. n = 40 trials. f. The mean correlation coefficient between imaging data and LFP recordings is significantly higher than that from shuffled control at each ROI (p < 0.001, Mann-Whiteney U test, n = 172 trials for both groups at each ROI). The control correlation coefficient is calculated from mismatched LFP recording and imaging trials. ROI locations are shown in (c). n = 3 mice, with 40-84 trials/mouse. g. LFP signals and JEDI-1P-Kv signals show correlation below 60 Hz based on the magnitude-squared coherence. Note that the dip in the coherence at 60 Hz is expected due to the LFP notch filter at this frequency.

We resolved the relation between LFP and JEDI-1P-Kv voltage at different frequencies using coherence analysis and found that coherence in anterior ROIs showed a clear peak around 8 Hz, which was stronger on the ipsilateral side. This peak was missing at posterior ROI locations (**Fig. 5e**). The analyzed site closest to the LFP recording showed coherence up to 60 Hz, while coherence with the ipsilateral motor cortex extended to 40 Hz. Coherence with contralateral structures was most dominant at frequencies below 8 Hz. These findings demonstrate the suitability of JEDI-1P-Kv widefield imaging to discover functional networks in different frequency bands up to 60 Hz, an important feature that has not been demonstrated for any other voltage indicators.

### JEDI-1P-Kv voltage signal tracks cortical voltage responses to whisker and visual stimulation trains up to 60 Hz

We next determined whether JEDI-1P-Kv’s fast kinetics would also enable the monitoring of fast neuronal responses to sensory stimulation under widefield imaging *in vivo*. We applied visual flicker from 20 Hz to 60 Hz to awake mice while imaging cortical activity (**Fig. 6a**). We determined which cortical areas showed voltage fluctuations following the stimulation frequency. Bilateral visual cortex produced the strongest response at the stimulation frequencies (**Fig. 6b**). To evaluate our ability to monitor how small cortical areas respond to different stimuli, we quantified JEDI-1P-Kv signals from a 2 × 2 pixel areas (200 × 200 μm) in the visual cortex. Single trial spectrograms showed a clear signal during the time of stimulation at the frequencies matching the ongoing visual flicker (**Fig. 6c**). Correspondingly, voltage traces during visual stimuli show oscillatory voltage fluctuations, indicating that populations of excitatory visual cortex neurons can follow high-frequency visual inputs (**Fig. 6d**).

**Fig 6.**
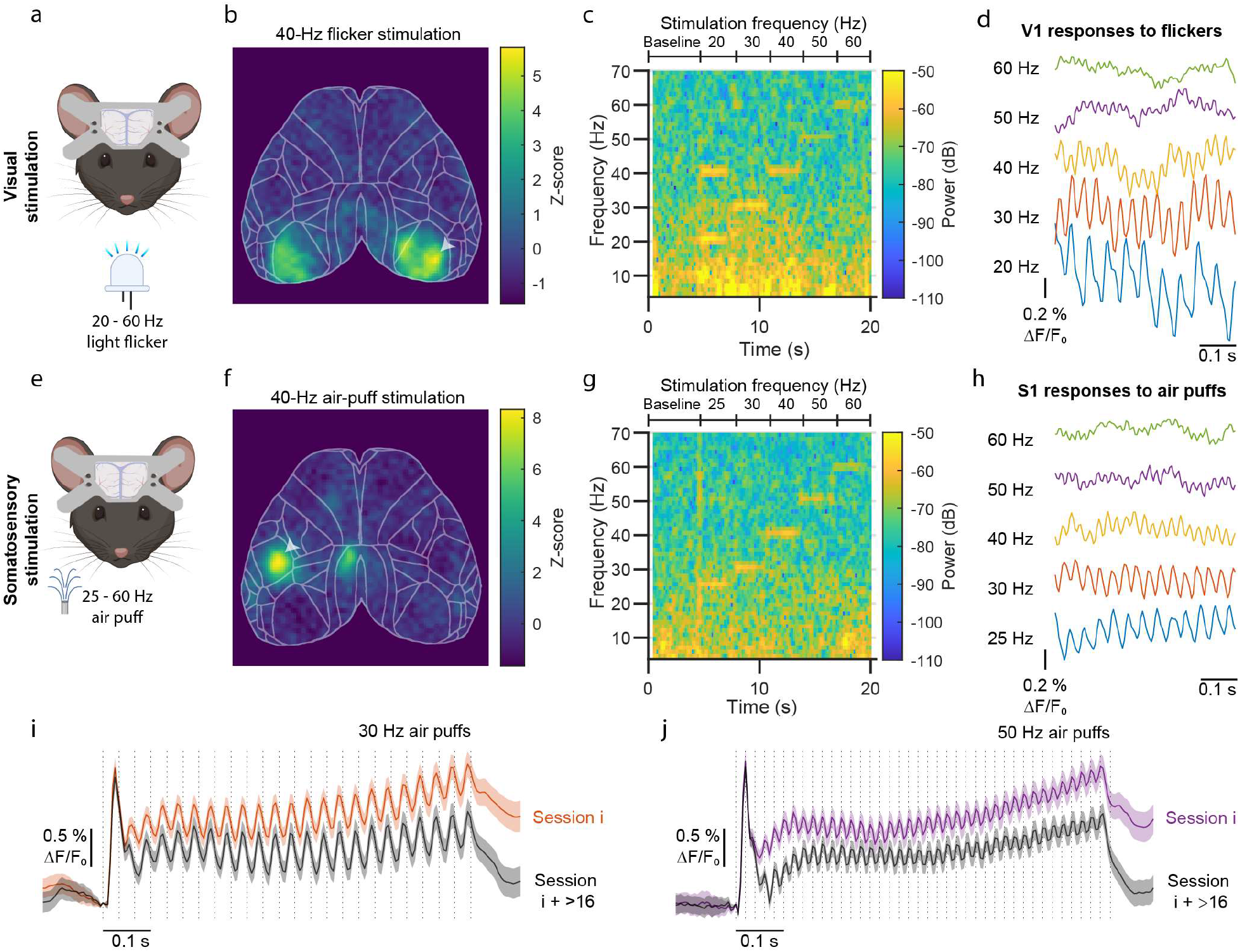
JEDI-1P-Kv reports rapid voltage responses to high-frequency stimuli. a-d. Simultaneous visual stimulation and widefield voltage imaging in awake mice. a. Experimental schematics. b. 40-Hz-filtered optical responses to 40-Hz visual stimulations are bilaterally localized in the visual cortex. The heat map shows the mean difference in the power spectral density at 40 Hz between stimulation and baseline from 10 trials. n = 1 mouse, representative of results with 4 mice. c. Single-trial imaging of JEDI-1P-Kv can report local voltage responses at frequencies matching visual stimulations of up to 60 Hz. Responses were acquired from a 2×2 pixel area in the visual cortex (arrow in f). n = 1 mouse, representative of results with 4 mice. d. Oscillations in the responses to specific flicker frequencies are visible in averaged voltage signals up to 60 Hz. Data from a 2×2 pixel area in the primary visual cortex. Data shown is from 1.5-2.0 s from the stimulus. n = 4 mice, 10 trials/mouse. e-h. Simultaneous whisker stimulation and widefield voltage imaging in awake mice. e. Experimental schematic. Air-puff trains were applied on the right side of whiskers. f. 40-Hz-filtered optical responses to 40-Hz air puffs are localized at the contralateral barrel cortex. The heat map shows the mean difference in the power spectral density at 40 Hz between stimulation and baseline from 10 trials. n = 1 mouse, representative of results with 5 mice. g. Single-trial imaging of JEDI-1P-Kv can report local voltage responses at frequencies matching air-puff stimulations of up to 60 Hz. Responses were acquired from a 2×2-pixel area in the contralateral barrel cortex (arrow in b). n = 1 mouse, representative of results with 5 mice. h. Same as d but for somatosensory stimuli and barrel cortex. Data shown is from 1.5-2.0 s from the stimulus. n = 5 mice, 10 trials/mouse. i-j. Responses to air puffs are stable over multiple days and imaging sessions. n = 216 trials from 6 mice (36 trials/mouse). Gray traces were acquired 16-19 imaging sessions after the orange and purple traces. The initial larger peak is consistent with electrophysiolocal data ^79^. Shaded areas denote the 95% CI.

To determine whether we could also detect fast voltage responses to somatosensory stimuli, we conducted widefield voltage imaging in response to 25-60-Hz air-puff trains (**Fig. 6e-h**). We filtered responses to monitor those matching the stimulus frequency and observed strong responses at the contralateral barrel cortex. Interestingly, we also detected robust responses in the contralateral retrosplenial cortex (RSPd) (**Fig. 6f**), an area most often associated with spatial navigation ^68^ and receiving some input from barrel cortex ^69^. Responses remained easily detectable over many recording sessions (**Fig. 6i,j**), supporting the use of JEDI-1P-Kv to study fast network dynamics during learning and other longitudinal approaches.

## Discussion

Neural computations underpinning behavior typically recruit distinct brain areas, motivating the need for large-scale and sensitive recording technologies. Here, we sought to address this critical need by engineering an improved GEVI and developing optimized widefield imaging techniques for pan-cortical voltage recordings with high spatiotemporal resolution.

To enable robust widefield voltage imaging, we developed and deployed an automated multi-parametric screening platform to evolve GEVI variants under widefield one-photon illumination. Our system can predict a variant’s brightness, photostability, response amplitudes to short and long voltage transients, and response width to spike waveforms. Our platform accelerated screening while avoiding outcomes where improvements in some characteristics are accompanied by decreases in others ^35^. The overall best indicator, JEDI-1P, is brighter and more photostable, and produces larger and faster responses to voltage transients (**Fig. 2**). These improved properties enabled 200-Hz voltage imaging through a cleared skull with high SNR (**Figs. 3d, 4a, 5a-b, 6**) and minimal photobleaching (**Fig. 3 f-h**). At ∼34°C, JEDI-1P has ∼0.5 and ∼1.2-ms on- and off-kinetics, respectively (**Table 1**), enabling single-trial reporting of voltage transients with similar timescales as LFPs (**Fig. 5b,e**) and neural responses to stimuli up to 60 Hz (**Fig. 6**).

We also reported improved approaches to promote rapid adoption of widefield voltage imaging by the neuroscience community. First, we demonstrated that soma-restricted JEDI-1P can be broadly and robustly expressed *in vivo* following ICV viral injections (**Fig. 3**), reducing costs by over an order of magnitude over retro-orbital injections. Compared with using transgenic animals, ICV viral injections enable rapid deployment of new GEVIs as they are developed. The components of our widefield imaging system (**Fig. 3b**) are commercially available, allowing end users to replicate our setup. The pre-processing pipeline we developed to correct for hemodynamic and motion artifacts is freely available (https://github.com/meowater/JEDI-1P-Kv_widefield_imaging_preprocessing_pipeline) and can be easily modified when used with an experimental preparation producing optical recordings with different noise characteristics (**Fig. 4**). While we applied JEDI-1P for pan-cortical widefield imaging, our new GEVI could be used for reporting neural activity using other methods including fiber photometry ^58^, miniaturized microscopes ^70^, confocal imaging ^71^, light sheet/SCAPE ^72^ and widefield imaging with cellular resolution ^73^.

Because JEDI-1P is exquisitely sensitive to subthreshold voltages (**Fig. 2a-b**) and cortical pyramidal neurons spend less than 1% of time spiking on average ^74^ a significant fraction of the signals reported by JEDI-1P-Kv under our conditions are likely hyperpolarizations and subthreshold depolarizations, which are not reported by calcium indicators. To our knowledge, by using JEDI-1P-Kv we provide the first method that can follow membrane voltage fluctuations at physiologically important frequencies of up to at least 60 Hz with long term stable imaging. Our new method therefore enables the cortex-wide study of beta and gamma oscillatory coupling in the dynamic control of functional networks as well as the relation between fast behavioral kinematics and their corresponding spatial neural response patterns.

Future work could combine supra-threshold measurements from calcium indicators with subthreshold recordings from JEDI-1P, either in separate animals, or by using a spectrally compatible calcium sensor ^75,76^. Because JEDI-1P is genetically encoded, follow-up studies could also explore the relative contributions of different cortical cell types – including interneurons ^77^ – to distributed neural computations. We look forward to the broader neuroscience community leveraging JEDI-1P and our optimized widefield imaging methods for understanding pan-cortical neural computations with cell type specificity and millisecond-timescale resolution.

**Figure S1.**
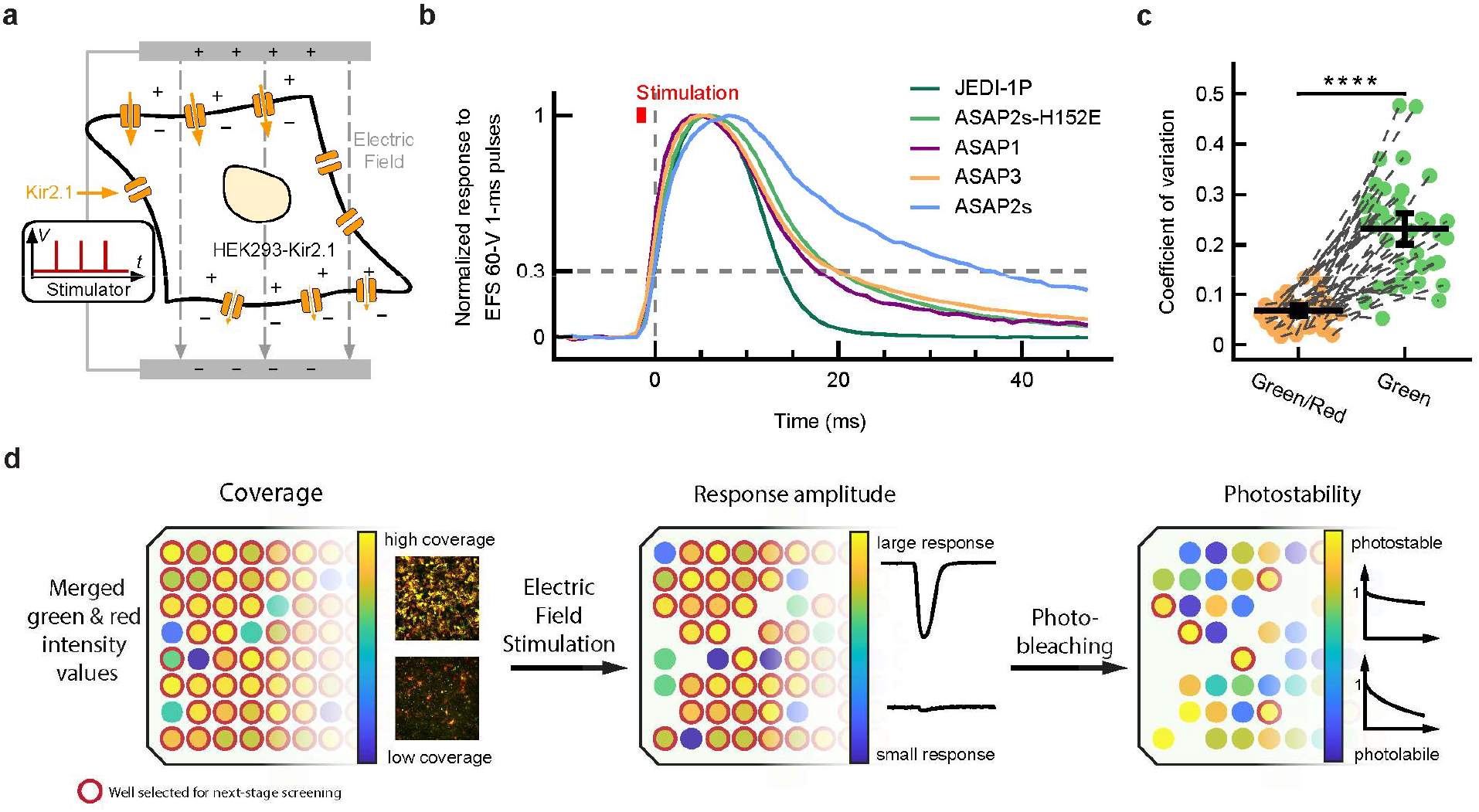
Design and benchmarking of the high-throughput multiparametric one-photon screening platform, related to Figure 1. a. Screening was conducted in HEK293 cells expressing Kir2.1 from the chromosome. Electric field stimulation (EFS) is thought to result in transient membrane hyperpolarization or depolarization, depending on the position of the membrane in the electrical field. Because Kir2.1 is inwardly rectifying, potassium inflows are larger than outflows, thereby causing net depolarization. b. 1-ms EFS produced transient depolarizations. Except for the slow indicator ASAP2s, other GEVIs reported these spikes with a fluorescence response lasting 10-20-ms measured at 30% of the peak response. c. To correct our quantification of brightness from well-to-well variations in overall expression level, we co-expressed GEVIs with the red fluorescent protein mCherry. Quantifying brightness as the ratio of GEVIs and mCherry produced a lower coefficient of variation than measuring brightness from GEVI fluorescence alone. **** p < 0.0001 paired t-test. Error bars denote 95% CI of the mean. d. Our screen evaluated various performance metrics in most wells. However, wells with the poorest expression typically produced noisy results; we thus excluded these wells from our response amplitude screen. To maximize screening throughput, we also did not evaluate photostability for constructs that did not respond to voltage.

**Figure S2.**
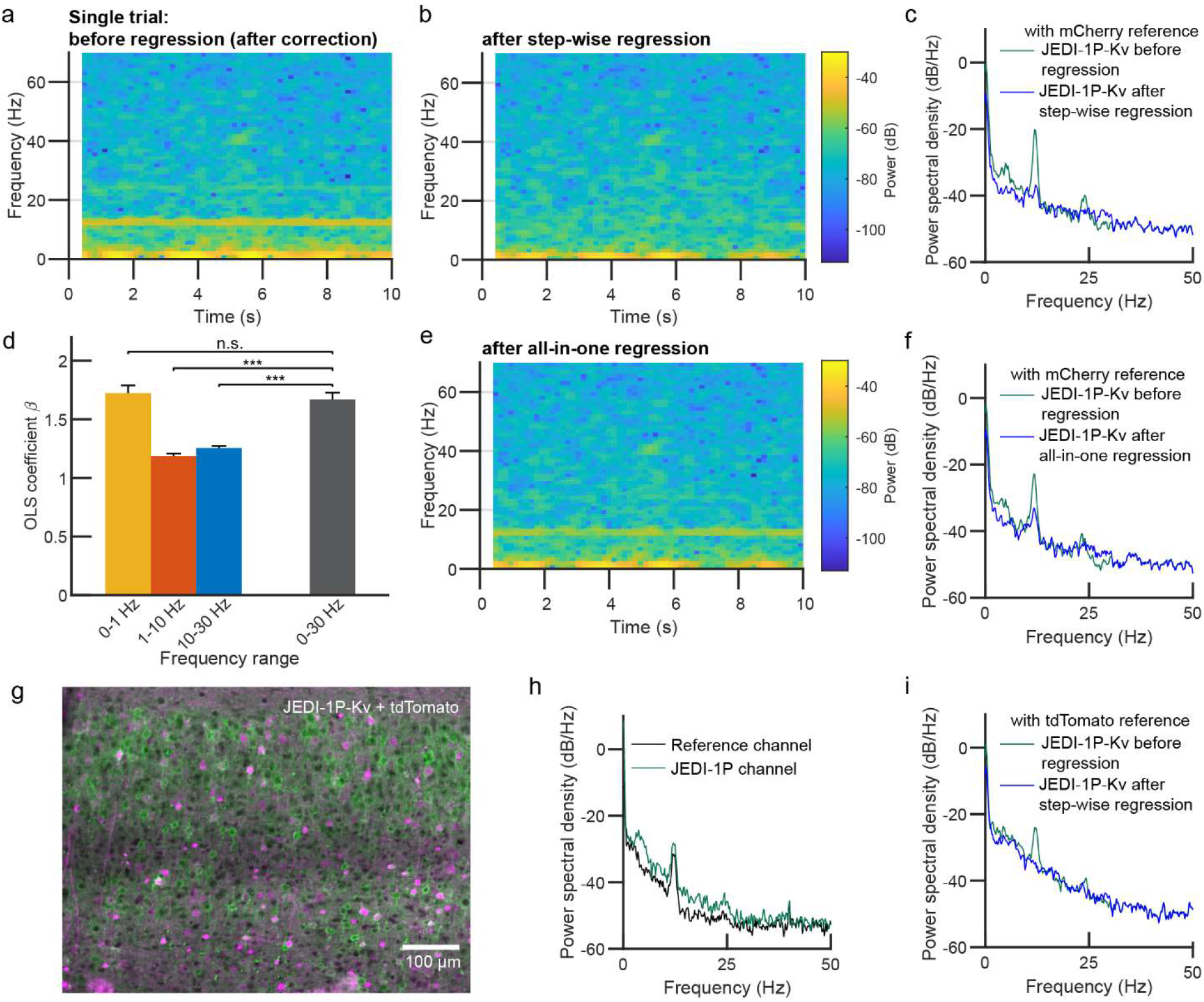
Regression is best done stepwise and can be achieved using different red fluorescent proteins. a-f. Conducting regression stepwise rather than all-in-one is more efficient at removing hemodynamic and motion artifacts. Data from awake mice. a-b. Single trial JEDI-1P-Kv spectrograms of neural activity from an awake behaving animal before (a) and after (b) regression illustrates that stepwise regression can remove fast hemodynamic noise. (a) and (b) share the same color bar. c. Most fast hemodynamic signals were removed from the voltage trace. n = 25 trials from a representative mouse. d. Noises from different frequency ranges have different amplitudes, which is reflected by significantly different scale factors (β) of Ordinary Least Square (OLS) regression. Error bars denote the 95% CI. n = 900 trials, 150 trials/mouse. ***, p < 0.0001; n.s., p > 0.05. Mann-Whitney U test. e. Computing all-in-one regression using a β value calculated from traces filtered with a broad frequency range retained significant artifacts at ∼12 and ∼24 Hz (yellow or high-dB horizontal bands). Same trial as (a) and (b), but with j = g - βr, where r was lowpass-filtered at 30 Hz. f. A representative all-in-step regression analysis retained significant fast hemodynamic noise, as shown by the significant peaks at ∼12 Hz and its harmonics at ∼24 Hz. n = 25 trials from the same mouse (same trials used in (c)). g-i. tdTomato can be used as an alternative reference protein to mCherry. g. Sagittal slice of the cortex shows expression of JEDI-1P-Kv and tdTomato. h. tdTomato can report hemodynamic noise, as shown by the clear peak at ∼12 Hz. Data is the mean of n = 20 trials at the same ROI from a representative mouse. i. The regression pipeline removed hemodynamic artifacts in mice co-injected with JEDI-1P-Kv and tdTomato AAVs. Data is the mean of n = 20 trials from a representative mouse.

## Methods

### High-throughput voltage indicator screening

#### Reagents and buffers

Cloning reagents include: PrimeSTAR HS DNA Polymerase (R040A, TaKaRa), FastDigest NheI (FD0974, Thermo Fisher Scientific), FastDigest HindIII (FD0504, Thermo Fisher Scientific), FastDigest KpnI (FD0524, Thermo Fisher Scientific), FastDigest Bsp1407I (FD0933, Thermo Fisher Scientific), FastDigest Green Buffer (10X) (B72, Thermo Fisher Scientific), agarose (BP160-500, Fisher BioReagents), Tris-Acetate-EDTA (TAE) 50× Solution (BP1332-1, Fisher BioReagents), Lysogeny broth, Miller (BP1426-500, Fisher BioReagents), Ampicillin (BP1760-25, Fisher BioReagents), In-Fusion HD Cloning Kits (639650, TaKaRa), GeneJET Gel extraction kit (FERK0691, Fisher), PureLink Pro Quick96 Plasmid Purification Kit (K211004A, Thermo Fisher Scientific) and Mini Plus Plasmid DNA Extraction System (GF2002, Viogene).

Cell and neuron culture reagents include: high-glucose Dulbecco’s Modified Eagle Medium (D1145, Sigma-Aldrich), fetal bovine serum (F2442, Sigma-Aldrich), glutamine (G7513, Sigma-Aldrich), Penicillin/Streptomycin (P4333, Sigma-Aldrich), Geneticin (G418) Sulfate (30-234-CR, Corning), phenol-free Neurobasal medium (12348017, Gibco), B-27 (17504044, Gibco), Glutamax (35050061, Gibco), 30-70 kD poly-D-lysine (P7886, Sigma-Aldrich), 300 kD poly-D-lysine hydrobromide (P7405, Sigma-Aldrich), Trypsin-EDTA solution (T3924, Sigma-Aldrich) and phosphate-buffered saline (PBS, SH302560, HyClone, GE Healthcare).

Transfection reagents include: jetPRIME (114-15, Polyplus Transfection), FuGENE HD transfection reagent (E2311, Promega), lipofectamine 2000 (11668019, Thermo Fisher Scientific) and Opti-MEM (31985-070, Gibco).

Reagents for solutions include: NaCl (S3014, Sigma-Aldrich), sucrose (S0389, Sigma-Aldrich), D-(+)-glucose (G8270, Sigma-Aldrich), HEPES (H3375, Sigma-Aldrich), KCl (P9541, Sigma-Aldrich), MgSO4 (M2643, Sigma-Aldrich), K-gluconate (P1847, Sigma-Aldrich), EGTA (E3889, Sigma-Aldrich), MgCl2 (M9272, Sigma-Aldrich), CaCl2 (223506, Sigma-Aldrich), KOH (P250, ThermoFisher) and NaOH (S5881, Sigma-Aldrich).

Used for culturing HEK293-Kir2.1 cells, **growth medium #1** contains high-glucose Dulbecco’s Modified Eagle Medium supplemented with 10% fetal bovine serum (FBS), 2 mM glutamine, 100 unit/mL Penicillin, 100 ug/mL Streptomycin, and 750 ug/mL of the antibiotic G418 Sulfate (geneticin).

Used in transfection and HEK293A cell culturing, **growth medium #2** contains high-glucose Dulbecco’s Modified Eagle Medium supplemented with 5% fetal bovine serum (FBS), 2 mM glutamine, 100 unit/mL Penicillin and 100 ug/mL Streptomycin.

Used in high-throughput screening, whole-cell patch-clamp, spectra characterization and neuron imaging, the **external solution** is composed of 110 mM NaCl, 26 mM sucrose, 23 mM glucose, 5 mM HEPES, 5 mM KCl, 2.5 mM CaCl2, 1.3 mM MgSO4 and is titrated to pH 7.4 with NaOH.

#### Plasmids construction and bacteria handling

All plasmids were constructed through standard molecular biology methods and sequence verified by Sanger sequencing. All primers for cloning were synthesized through Sigma-Aldrich standard DNA oligos service.

For vectors used for screening, GEVIs were fused to mCherry-CAAX at the C-terminus ^47,81,82^ via a self-skipping P2A linker ^83^ and were cloned in pcDNA3.1/Puro-CAG vector. ASAP1 and ASAP2s were subcloned from Addgene plasmids #52519 and #101274, respectively. ASAP3 was subcloned from a plasmid kindly provided by Dr. Michael Lin (Stanford). ASAP1-EGFP was cloned by replacing the circularly permuted GFP in ASAP1 (cpsfGFP-OPT) with EGFP (V2 – K239 ^35^(13)). ASAP1-dpOPT was cloned by replacing the cpsfGFP-OPT in ASAP1 with de-permuted cpsfGFP-OPT (dpsfGFP-OPT).

Polymerase chain reaction (PCR) was used to create single-site-saturation mutagenesis libraries. To achieve a uniform distribution of the amino acid, we mix a set of 4 forward primers at a molar ratio of 16:3:1:1, each containing a (degenerate) codon at the mutagenesis site: NNT (N = A, T, G or C), VAA (V = A, G, or C), ATG, or TGG. For each PCR reaction of 20 μL, the mixture of the forward primers and the reverse primer were each added to a final molar concentration of 0.5 μmol, together with 10-50 ng template plasmid and 10 μL 2×PrimeSTAR DNA polymerase premix. DNA was amplified using the following protocol: an initial denaturation step at 98°C for 30 s; 35 amplification cycles of 98°C for 10 s, 57°C for 10 s, 72°C for 1 min/kb of fragment length; a final extension step at 72°C for 5 min. The backbone was linearized from pcDNA3.1/Puro-CAG vector with a ccdB-camR insert, using restriction enzymes NheI and HindIII. PCR products and linearized vector were both purified with gel electrophoresis and GeneJET Gel extraction and assembled via In-Fusion seamless cloning strategy per manufacturer’s instructions. 1 uL In-Fusion reaction mix was transformed into commercial chemically competent *Escherichia coli* (XL10-Gold, Agilent). Colonies were manually picked into Lysogeny broth supplemented with 100 μg/mL ampicillin in a 96-well block the next day after transformation and incubated in a shaker (MaxQ 5000, Thermo Scientific) kept at 37°C, and plasmids were purified 18 h after the inoculation with a 96-well plasmid purification kit (PureLink Pro, Thermo Fisher Scientific) following the manufacturer’s instructions.

For vectors used for *in vitro* characterization in HEK293A cells, GEVIs were cloned in pcDNA3.1/Puro-CAG vectors with the same cloning strategy as constructing screening vectors except for no reference protein was attached, no degenerate codons were used, and the plasmids were purified with Viogene mini plus miniprep kit.

For vectors used for one-photon excitation and emission spectra characterization in HEK293-Kir2.1 cells, a control vector pcDNA3.1/Puro-CAG-EGFP-CAAX was constructed by fusing the CAAX membrane anchoring motif to the C-terminal of EGFP. Same cloning and DNA preparation strategies were used as the vectors used for in vitro characterization.

For vectors used for dissociated neuron expression, JEDI-1P was cloned into pAAV vector under the control of the neuron-specific hSyn promoter by replacing the ASAP2s sequence in pAAV-hSyn-ASAP2s (Addgene plasmid #101276). pAAV-hSyn vector was linearized with restriction enzymes KpnI and HindIII.

For vectors used for mice expression, we attached a Kv2.1 PRC motif to the C-terminus of JEDI-1P through a triple GSS linker (GSSGSSGSS) to restrict the expression to soma and proximal dendrites. The double foxed inversed JEDI-1P-Kv2.1prc cassette was cloned into the pAAV vector under the control of EF1α promoter by replacing the hChR2(H134R)-EYFP sequence in pAAV-EF1α -double floxed-hChR2-EYFP-WPRE-HGHpA (Addgene plasmid #20298). pAAV-EF1a vector was linearized with restriction enzymes Bsp1407I and NheI.

#### Mammalian cell culture and transfection

For GEVI screening, we use a modified HEK293 cell line that stably expresses human Kir2.1 channel to maintain a resting membrane potential of around −77 mV ^44^(20). Cells were cultured in Sanyo CO2 incubators (MCO-18AIC(UV), Marshall scientific) that are kept at 37°C and supplemented with 5% CO2. **Growth medium #1** that contains 750 ug/mL of the antibiotic G418 Sulfate was used to maintain Kir2.1 expression in the cell line.

Transfections of HEK293-Kir2.1 cells for screening were performed in 96-well glass-bottomed plates (P96-1.5H-N, Cellvis). Plates were coated with 30-70 kD poly-D-lysine for 1 h at 37°C to promote cell adhesion and washed twice with PBS before seeding the cells. HEK293-Kir2.1 cells were then seeded one day before forward transfection at 60-65% confluency, equivalent to 24,000-26,000 cells per well, or 1 h before reverse transfection at 70-80% confluency, equivalent to 28,000-32,000 cells per well. **Growth medium #2** that is supplemented with 5% FBS and free from G418 was used for seeding the cells to improve the cellular health while restricting the growth rate. Transfections were performed 36-48 h before screening with JetPRIME according to their protocol: for each well of the 96-well plate, 0.4 μL of JetPRIME reagent was first mixed in 10 μL of JetPRIME buffer, and then mixed with 130 ng of DNA prediluted in 10 μL JetPRIME buffer. The mixture was incubated for 8 min at room temperature before adding to the wells seeded with cells, and the medium in the 96-well plate was replaced with fresh **growth medium #2** the next day to minimize the cytotoxicity of the transfection reagent.

#### High-throughput GEVI screening platform

To evaluate the performance of GEVIs in a high throughput way, we built an automated multimodal 96-well screening platform based on an inverted microscope (A1R-MP, Nikon Instruments). The excitation light was generated from an LED light engine (SpectraX, Lumencor) and directed to the microscope’s epifluorescence illuminator via a liquid light guide. Cyan (470/24 nm, center wavelength/bandwidth) and Green (550/15 nm) light were delivered to the sample plan through a 20× 0.75 NA objective (CFI Plan Apochromat Lambda, Nikon Instruments) to excite the GFP-based GEVIs and mCherry, respectively. The emission light from the sample plane was split from the excitation light through a multi-band dichroic mirror (89000bs, Chroma) and filtered by a multi-band emission filter (89101m, Chroma) before being collected by a scientific complementary metal–oxide–semiconductor (sCMOS) camera (ORCA Flash 4.0 V2, C11440-22CU, Hamamatsu). A motorized extended travel stage capable (H139E1, Prior) was used to control the position of the field of view and to hold 96-well plates.

To support automation of the system, data acquisition and output broads (PCI-6229 and PCI-6723, National Instruments) were connected to the microscope computer through a PXI Chassis (PXI-1033, National Instruments). The computer was equipped with 2 Intel Xeon E5-2630 v3 processors (total of 16 cores), 128 GB of DDR4 RAM, and four 2 TB SSDs in RAID 0 to facilitate high-speed imaging. JOBs scripts in NIS-Elements HC (version 4.60, Nikon Instruments) were used to control the microscope system (e.g., stage position), manage the optical configurations (e.g., excitations), initiate image acquisition, and trigger the stimulator.

A digital isolated high-power stimulator (4100, A-M System) was used to provide electric field stimulation. Electric pulses were passed to a pair of electrodes made from 0.5 mm wide platinum wires (99.95% pure, AA10286BU, Fisher Scientific). The two L-shaped electrodes had a horizontal length of 2 mm and were 3 mm apart (**Fig. 1b**), and they were secured on a 3D-printed polylactic acid holder. The holder was fixed to a motorized linear translation stage (MTS50-Z8, Thorlabs), which was controlled by Kinesis (Thorlabs) to move the electrodes in and out of individual wells. Two smaller manual linear translation stages (411-05S, Newport) were used to fine-tune the electrodes’ lateral position. During stimulation, the electrodes were submerged under the imaging solution, about 500 μm above the bottom.

#### High-throughput GEVI screening under widefield one-photon illumination

On the day of screening, medium in the 96 well plate was removed and cells in each well were washed with 200 μL **external solution** twice, before another 100 μL of **external solution** was added to each well as the imaging solution. To determine the brightness and the response amplitude of the sensors, 4 non-overlapping fields of view (FOVs) of 2048 × 200 (512 × 50 post 4 × 4 binning) pixels were imaged per well, allowing the sCMOS camera to record at a maximum speed of 987 frames per second (fps). The electric field stimulation (EFS) electrode was moved into the well after the motorized stage centered at the well and before the stage moved to the first selected FOV. The reference channel designated for the brightness of the mCherry was captured first under 555 nm illumination for 1 frame, with an irradiance of ∼69 mW/mm^2^ at the sample plane and an exposure time of 5 ms. Then, the target channel designated for capturing the responses from the green GEVIs was imaged at ∼987 fps under 470 nm illumination for 4900 frames (∼5 s), with an irradiance of ∼36 mW/mm^2^ at the sample plane and an averaged exposure time of 1.01 ms. Immediately before the camera started, a transistor–transistor logic (TTL) signal was sent to trigger the stimulator, which sent out a TTL signal to trigger the shutter of the light source and turn light on. This allowed the camera to capture the onset of the excitation light, which eventually served as a reference point to locate the EFS events and temporally align the results from different FOVs. The stimulations took place 1 s after the light was turned on, started from four monophasic square pulses with 1-ms width, 60-V amplitude, and a period of 500 ms. This was followed by a 100-Hz train stimulation (10 monophasic square pulses with 2.5-ms width, 30-V amplitude, and a period of 10 ms), with a 1-s gap between the single pulses and the train stimulation. After the EFS on the 4 selected FOVs, the motorized stage centered in the well again, and the motorized linear translation stage raised the electrode outside of the well.

To determine the photostability of the sensors in each of the well, another FOV of 2048 × 2044 (512 × 511 post 4 × 4 binning) pixels away from the previous FOVs was bleached for 2 min under 470 nm with an irradiance of ∼36 mW/mm^2^ at the sample plane, and the photobleaching video was recorded at a frame rate of 100 fps and an averaged exposure time of 10 ms. The excitation light was triggered the same way as described above (but no electrode in the well so no EFS took place) to generate the light onset signal for aligning the results from different FOVs. Before each photobleaching video was taken, another single-frame reference channel for evaluating the mCherry intensity in the same FOV was captured with 555 nm LED excitation, with an irradiance of ∼69 mW/mm^2^ at the sample plane and an exposure time of 5 ms.

#### High-throughput screening data analysis

Time-lapse images collected from the screening were processed by custom routines in MATLAB (version r2019b, MathWorks). Recording files in Nikon NIS-Elements ND2 format were first parsed by the Bio-Formats MATLAB toolbox (version 6.3.1) ^84(47)^. For both channels of each FOV, saturated pixels, typically from over-expressing cells, were excluded and background levels were estimated using the intensity histogram and subtracted from images. To exclude pixels that do not represent cells, an initial mask was computed from the first frames of each channel using predefined thresholds. This mask was applied to the entire recording, and the overall change of fluorescence over time can be obtained by summing pixels selected by the mask frame by frame across all time points.

For electric field stimulation recordings, further pixel selection was necessary to mitigate pernicious influences on response amplitude due to overexpressing cells, bright extracellular fluorescent puncta, and intracellular aggregates. Based on the initial mask, for the green channel time series, a secondary mask with non-responsive pixels removed was generated to obtain response amplitudes. To do this, we first estimated the photobleaching by fitting a three-term exponential curve on the overall fluorescence with time points during stimulation excluded. The time constants from the fitting were then used to estimate the trend for each pixel using the least-squares fitting. The trend of each pixel was then removed using division. After detrending, we calculated correlation between the overall fluorescence to that of each pixel within the initial mask. These pixels were then ranked with decreasing correlation and grouped in batches of 200 pixels. The secondary mask was determined by aggregating groups of pixels from the highest correlation until the signal-to-noise ratio reached a maximum. Fluorescence from the secondary mask was summed up and normalized by initial fluorescence for each FOV to quantify response amplitude.

For photobleaching recordings, average fluorescence from pixels in the initial mask was normalized by the initial fluorescence. To quantify brightness, pixels of each channel from the initial mask were first averaged and then the brightness was defined as the ratio of mean green fluorescence over red fluorescence.

### *In vitro* characterization of JEDI-1P

#### Reagents and buffers

Used for plating dissociated neurons, the **neuron plating medium** contains Neurobasal medium supplemented with B-27, 2 mM Glutamax, 10% FBS, 100 unit/mL Penicillin, and 100 ug/mL Streptomycin.

Used for culturing dissociated neurons, the **neuron culturing medium** contains phenol-free Neurobasal medium supplemented with B-27, 2 mM Glutamax, 100 unit/mL Penicillin, and 100 ug/mL Streptomycin.

Used in whole-cell patch-clamp, the **internal solution** is composed of 115 mM K-gluconate, 10 mM HEPES, 10 mM EGTA, 10 mM glucose, 8 mM KCl, 5 mM MgCl_2_, 1 mM CaCl_2_ and is adjusted to pH 7.4 with KOH.

#### Mammalian cell culture and transfection

For *in vitro* GEVI characterization with whole-cell voltage clamp, we used HEK293A cells (R70507, Thermo Fisher scientific). The cell line was cultured under the same condition as HEK293-Kir2.1 cells except that **growth medium #2** was used with no G418 Sulfate supplement.

Transfections of HEK293A cells for patch clamp were performed in 24-well plates (P24-1.5H-N, Cellvis) on 12-mm cover glass (#0, 633009, Carolina). Cover glass was coated with 30-70 kD poly-D-lysine for 5-10 min at 37°C to promote cell adhesion and washed twice with PBS before seeding the cells. HEK293A cells were seeded in **growth medium #2** two days before patch clamp at 30% confluency, equivalent to 72,000-96,000 cells per well. Transfections were performed on the same day of seeding per manufacturer’s instruction: for each well of the 24-well plate, 0.6 μL FuGENE transfection reagent that has pre-equilibrated in room temperature for 15 min was added to 200 ng DNA diluted in 13 μL Opti-MEM. The mixture was incubated for 7 min at room temperature before adding to the cells, and the medium was replaced with fresh **growth medium #2** the next day.

#### Whole-cell voltage clamp setup

To evaluate the performance of GEVIs with whole-cell voltage clamp, we used GEVIs with no reference protein attached cloned in pcDNA3.1/Puro-CAG vectors and transfected into HEK293A cells as described in **Cell culture and transfection**. Electrophysiological recordings were done 36-48 h post transfection at room temperature unless otherwise stated.

Pipettes for patch clamping were prepared freshly on the day of electrophysiology experiments from filamented glass capillaries (1B150F-4, World Precision Instruments) using a horizontal pipette puller (P1000, Sutter) to achieve a tip pipette resistance of 3-5 MΩ. The micropipettes filled with the **internal solution** using MicroFil Flexible Needle (MF28G67-5, World Precision Instruments) were secured on a patch-clamp headstage (CV-7B, Molecular Devices) and positioned by a micromanipulator (SMX series, Sensapex).

The coverslip seeded with the transfected cells was placed in a custom glass-bottom chamber based on Chamlide EC (Live Cell Instrument) with glass bottom made with a 24 × 24 mm #1 coverslip (89082-270, VWR). Cells were continuously perfused with the **external solution** at ∼ 4 mL/min with a peristaltic pump (505DU, Watson Marlow). Whole cell voltage clamp was achieved using a MultiClamp 700B amplifier (Molecular Devices). Patch clamp data was recorded with an Axon Digidata 1550B1 Low Noise system with HumSilencer (Molecular Devices). Cells were held at −70 mV at the baseline. A liquid junction potential of 11 mV was compensated in all command voltage waveforms. Recordings were considered satisfactory and were included in the final analysis only if the patched cell had an access resistance (Ra) smaller than 12 MΩ and a membrane resistance (Rm) larger than 10 times Ra both before and after the recording.

All patch clamp recordings were performed on the same inverted microscope used for GEVI screening. Cyan (470/24 nm, center wavelength/bandwidth) excitation light generated from the SpectraX light engine was delivered to the sample plane through a 40× 1.30 NA objective (CFI Plan Apochromat Lambda, Nikon Instruments) to excite the GEVIs at an irradiance of ∼ 37.41 mW/mm^2^ unless otherwise noted.

#### Evaluating indicators’ response amplitude

To evaluate the sensors’ response to action potentials (APs), we voltage-clamped the cells to follow a typical AP waveform that was recorded from a representative hippocampal neuron and modified to have an amplitude of 100 mV (from −70 mV baseline voltage) and a full width at half maximum of 2 ms to mimic the shape of layer 2/3 cortical neurons at room temperature ^85^. Cells were stimulated with 5 single AP waveforms at 2 Hz and 10 AP waveforms in train at 100 Hz, with around 3 s of interval holding at −70 mV between the two protocols. The voltage-clamped cell was centered in a FOV of 2048 × 200 (512 × 50 post 4 × 4 binning) pixels. The emission light from the cell was filtered by a multi-band emission filter (89101m, Chroma) and collected by the ORCA Flash 4.0 V2 sCMOS camera at ∼987 Hz with an averaged exposure time of 1.01 ms.

To evaluate the sensors’ response to steady-state voltages, we applied 1-s step-voltage stimulations of −100, −80, −60, −40, −20, 0, 20, 30, and 50 mV to the voltage-clamped cells, with a 1.5 s gap holding at −70 mV between each two stimulations. The emission light was captured with the same optical configuration and camera settings as AP capturing as described above.

Time series recordings from the camera were processed using a similar method as for processing high-throughput electric field stimulation recordings (see above).

#### Evaluating indicators’ kinetics

To evaluate the sensors’ kinetics, we applied three 1-s 100-mV depolarization pulses from −70 to 30 mV. Between two pulses, cells were held at −70 mV for 1.4 s. Electrophysiological recordings were conducted at 21-23°C (room temperature) or 32-35°C (closer to the 37°C temperature of mice brains) using a feedback-controlled inline heater system (inline heater SH-27B, controller TC-324C, cable with thermistor TA-29, Warner instruments) to maintain the temperature in the perfusion chamber. Irradiance of the excitation light at the sample plane was tuned down to ∼19.43 mW/mm^2^ to minimize photobleaching, and a diaphragm was used to reduce the diameter of the excitation spot to ensure only the patch-clamped cell was illuminated.

To capture fluorescence changes at a higher temporal resolution than that of our camera, we used a multialkali photomultiplier tube (PMT, PMM02, Thorlabs) installed on one of the side ports of the microscope to collect the emitted photons from the GEVIs. A LabVIEW (version NXG 5.0, National Instruments) routine was used to control the PMT bias voltage and record the output voltage using the data acquisition and output boards. Data was collected at 80 kHz.

The output voltage from the PMT was analyzed by a custom routine written in MATLAB to obtain fluorescence signal for each cell. The raw data was first downsampled to 20 kHz. Then, photobleaching correction was done by performing a three-term exponential fitting on the baseline (when the cell was held at −70 mV) and removing the trend from the entire signal using division. The corrected signal was cropped from 0.1-s before the estimated depolarization or repolarization onset to 1-s after the estimated depolarization or repolarization onset. The exact onset timing was fitted together with other coefficients with either single-exponential (F(t) = c + (k × exp((t – t_0_) × λ)) × (t > t_0_) + k × (t <= t_0_)) or dual-exponential ((F(t) = c + (k × exp((t – t_0_) × λ) + k_2_ × exp((t – t_0_) × λ_2_)) × (t > t_0_) + (k + k_2_) × (t <= t_0_)) model where the t is the independent variable, F is the dependent variable, and the rest are the coefficients to be fitted. Among these coefficients, c describes the mean plateau fluorescence, k and k2 describe the relative ratio of each exponential component, λ and λ_2_ describe the opposite reciprocal of time constants, and t0 is an offset indicating the exact event onset timing.

#### One-photon excitation and emission spectra characterization

To characterize the one-photon excitation and emission spectra of JEDI-1P, we used the same vector used for *in vitro* GEVI characterization with whole-cell voltage clamp, i.e., pcDNA3.1/Puro-CAG-JEDI-1P, and a control vector pcDNA3.1/Puro-CAG-EGFP-CAAX expressing membrane-anchored EGFP to mimic the spatial localization of JEDI-1P within the cells. HEK293-Kir2.1 cells were used to express JEDI-1P or EGFP-CAAX to characterize the spectra of the indicator at a membrane voltage that is close to the resting membrane potential (–70 mV). Transfections were performed in 6-well plastic plates (3516, Corning), and HEK293-Kir2.1 cells were seeded one day before forward transfection at 60-65% confluency, equivalent to 720,000-780,000 cells per well in **growth medium #2**. Transfections were performed 36-48 h before imaging using 9 μL JetPRIME and 200 μL JetPRIME buffer per well. The transfection medium was replaced 4 h after transfection with fresh **growth medium #2** to minimize potential cytotoxicity from the transfection reagent.

On the day of experiment, cells from 2 wells of the 6-well plate transfected with the same construct were detached with trypsin, washed twice with and diluted into the same **external solution** for screening and *in vitro* characterization, and pooled into a single well of a glass-bottomed 96-well plate (P96-1.5H-N, Cellvis). Pooling the cells to a dense preparation was important to produce a strong signal that could be robustly detected by the plate reader. Untransfected cells were prepared with the same method to determine the background autofluorescence levels. A hemocytometer was used to plate a similar number of cells between conditions.

Spectra were determined by using a plate reader (Cytation 5, BioTek) to quantify fluorescence from wells of the 96-well plates prepared above. Excitation spectra were acquired by scanning excitation wavelengths from 350 to 535 nm in increments of 1 nm and a bandwidth of 10 nm and collecting emission intensity at 560/10 nm. Emission spectra were acquired by exciting at 430/10 nm and measuring emitted photons from 460 to 650 nm in increments of 1 nm and a bandwidth of 10 nm. Individual scans of excitation and emission spectra were corrected for autofluorescence by subtracting the values from untransfected cells at each wavelength, and then normalized to their respective peaks. The final excitation and emission spectra were averaged from normalized spectra of each individual scan. The peaks were averaged from the peaks of each individual scan.

#### Confocal imaging of GEVI in dissociated neurons

To determine the expression and trafficking of GEVIs in neurons, we used the rat cortical neurons. Primary rat cortical neurons were isolated from day 18 Long-Evans rat embryos. Cortices were dissected, dissociated with papain (Worthington Biochemical Corporation), washed with trypsin inhibitor (Sigma), and seeded at 500,000 cells/mL in 500 μL **neuron plating medium** per well of a 24-well glass bottom plate (P24-1.5H-N, Cellvis). The plate was pre-coated overnight with 300 kD poly-D-lysine hydrobromide and washed twice with PBS before seeding. The plating day was considered as day in vitro (DIV) 0. The next day, 90% of the media was replaced with the **neuron culturing medium**. Half of the media was henceforth replaced with fresh **neuron culturing medium** every 3-4 days. All media were pre-equilibrated for at least 24 h at 37°C in air with 5% CO_2_ before usage. Neurons were kept in Sanyo incubators (MCO-18AIC(UV), Marshall scientific) that are kept at 37°C and supplemented with 5% CO_2_ after plating, and transfected at DIV 9 using 1 μL lipofectamine 2000 and 800 ng total DNA, including 200 ng pAAV-hSyn-JEDI-1P and 600 ng pNCS bacterial expression vector as buffer/filler DNA.

Six days after transfection (DIV 15), the attached neurons were washed twice with the **external solution** before adding a final 500 μL per well as the imaging solution. Laser-scanning confocal images were obtained using a high-speed confocal microscope (LSM880 with Airyscan, Zeiss) driven by the Zen software (version 2.3 SP1, Zeiss). The microscope was equipped with a 40× 1.1-NA water immersion objective (LD C-Apochromat Korr M27, Zeiss), a 488-nm argon laser (LGK7812, Lasos) set to 20% power (∼200 μW) and a per-pixel dwell time of 2.35 μs. Emission light was filtered using a multipass beamsplitter (MBS 488/561/633, Zeiss) and acquired with a 32 channel GaAsP detector (Airyscan, Zeiss) with a detector gain of 800, and a 2.15-Airy unit pinhole size. Images were acquired at a resolution of 0.05 μm/pixel and an X-Y dimension of 3544 × 3544 pixels. Z stacks were stepped at 0.22 μm between images. **Fig. 2k** corresponds to a maximum intensity projection from Z-stack with 25 images. Airyscan processing was applied to the images to increase the resolution.

### Widefield imaging of JEDI-1P in mouse

#### Animals

14 male and 15 female *EMX1*-Cre mice were used for experiments. Animals were provided with *ad libitum* food and water unless specified otherwise. For imaging experiments, mice that received imaging window and head plate implantation surgery were singly housed with enrichment under a reverse cycle (12 h light, 12 h dark). Imaging experiments were performed during the dark cycle. Mice were placed on continuous water restriction for awake imaging. Emory University Institutional Animal Care and Use Committee approved all animal work performed in this study.

#### Neonatal intracerebroventricular injection

To achieve brain-wide expression of JEDI-1P and reference fluorescence, we used intracerebroventricular injection to deliver viral vectors ^61^. *EMX1*-Cre mice were used to set up mating pairs. Breeder cages were monitored twice daily approaching the expected delivery day. Newborn pups were carefully retrieved from the breeder cage after they showed visible milk spot. The pups were then placed on a heating pad kept at 37°C during the preparation of viral vectors. AAV.php.eb-EF1a-DIO-JEDI-1P-Kv2.1-WPRE (2-4 × 10^12^ vg/mL) and AAV9-hSyn-mCherry (3.1 × 10^13^ vg/mL) were mixed with a particle ratio of 1:3. For mice that received tdTomato instead of mCherry, the viral solution was consisted of AAV9-EF1a-DIO-JEDI-1P-Kv2.1-WPRE (1.9 × 10^13^ vg/mL) and AAV.PHP.eb-CAG-tdTomato (3.1 × 10^13^ vg/mL) with a particle ratio of 38:1. For injection in each pup, a total of 5 μL mixture was loaded to a 10-μL Nanofil syringe (World Precision Instrument) with a 34G beveled needle (NF34BV-2, World Precision Instrument). After identifying the injection site, 2/5 of the distance between the lambda suture to each eye, the syringe was carefully inserted at the target to a depth of 3 mm. 2 μL of viral vectors were slowly dispensed on each hemisphere. After all the injections were complete, the pups were returned to the breeder cage.

#### Histology

Mice were injected with 0.1 mL of Euthasol (51311-0050-01, Virbac) prior to transcardial perfusion. The animals were then perfused with 0.1 M phosphate buffer (PB) followed by 4% paraformaldehyde (PFA) PB solution. The harvested brain tissue was left in 4% PFA PB solution for 24 h before being transferred to 15% sucrose PB and then 30% sucrose PB. The brain tissue was sectioned with a microtome (HM430, Epredia) at 50 μm thickness and imaged with a fluorescence microscope (BZ-X, KEYENCE).

#### Surgery of imaging window and head-post implant

Imaging window for *in vivo* widefield imaging was adapted from the clear-skull method ^62,63^(38,39) 3% isoflurane was used for induction of anesthesia, and mice were kept under 2% isoflurane during the surgery. After fur removal with Nair (I0041395, Church & Dwight), the scalp was cleaned with alternating 70% ethanol and povidone-iodine pads. The scalp was removed to expose about 9 × 9 mm of the skull. The skull was gently scraped with a scalpel to remove fascia and dried with cotton swabs. A thin layer of Opti-bond Universal (36519, Kerr) was then applied to the dried surface. After Opti-bond was cured with ultraviolet light, a layer of dental cement mixture was applied before placing the custom cut cover slip (12-545-88, Fisherbrand Superslip) in place. The cover slip was cut with a diamond wedge scribe (S90W, Thorlabs) to a hexagon shape that was fitted to the inside of a head-plate (**Fig 3c**). The dental cement for creating clear imaging window was mixed with 1 scoop of C&B Metabond L-powder and 5 drops (S399, Parkell Products Inc.) of C&B Metabond Quick Base (S398, Parkell Products Inc.). Head plate for head-fixation was also secured with the dental cement (2 scoops of L-powder and 6 drops of Quick Base) after the imaging window was cured in place.

#### Continuous water restriction

For awake imaging sessions, mice were placed under water restriction at least 2 days prior to the experiments. On day one of the water restriction, the ad libitum water was removed. The weight of the animal at the time of water removal was recorded as the baseline weight. The animal was weighed daily and was given a minimum of 40 ml/kg water each day (according to its baseline weight) to maintain a weight above 80% of the baseline. Continuous water restriction lasted no longer than 2 months.

#### In vivo widefield imaging

A CMOS high-speed imaging system (MiCAM ULTIMA. SciMedia Ltd.) with dual cameras was used for *in vivo* widefield imaging at 200 Hz (**Fig. 3b**). JEDI-1P-Kv and mCherry were excited with a single blue LED (466/40 nm, center wavelength/bandwidth, FF01-466/40, Semrock) through the imaging window. The power of excitation light was 0.05 mW/mm^2^ at the plane of imaging window for all experiments. Emission light from JEDI-1P-Kv and mCherry was conditioned by 525/50 nm (Semrock: FF03-525/50-50) and 650/60 nm (FF01-650/60-50, Semrock) emission filters, respectively. For imaging of mice with JEDI-1P-Kv and tdTomato, the emission filter for JEDI-1P-Kv was replaced with 515/30 nm (FF03-515/30-50, Semrock) to avoid emission from tdTomato above 530 nm. A dichroic beam splitter at 495 nm (FF495-Di03-50×70, Semrock) was used to reflect excitation light, and another dichroic at 580 nm (FF580-FDi01-50×70, Semrock) was used to split the emission light into two cameras. The spatial resolution of each camera was 100 × 100 pixels, and the field of view was 10 × 10 mm in size.

#### Electrocardiography recording

To qualitatively verify that fast hemodynamic signals are present in the imaging signals, we recorded electrocardiography (EKG) simultaneously during imaging. The fur on the animals’ chest was shaved on the day of recording. Mice were lightly anesthetized with 1% isoflurane during EKG recording. A circular shaped electrode with Spectra 360 electrode gel (12-08, Parker Labs) was secured onto the shaved chest area. The EKG signals were amplified through Extracellular Amplifier (EXT-02F, NPI Electronic GmbH) and recorded simultaneously during imaging.

#### Widefield imaging data processing

We developed a pipeline that uses reference fluorescence to subtract non-voltage signals from JEDI-1P-Kv signal. We implemented the data processing algorithms in MATLAB (version R2021b). The background fluorescence, estimated by measuring autofluorescence from control mice (n = 4 mice) under the same imaging condition, was first subtracted from both JEDI-1P-Kv channel and reference channel. Then the photobleaching was detrended from the background subtracted fluorescence JEDI-1P-Kv trace. The fast photobleaching was calculated empirically from averaging 5536 JEDI-1P-Kv traces (20.48 s long, n = 6 mice, 692 trials, 8 traces/trial), which were digitally filtered with a lowpass filter (*lowpass* function) at 0.5 Hz. The mean fast photobleaching was then fitted to a two-term exponential model. We also employed the same process to estimate the photobleaching from mCherry but found no apparent photobleaching in the mCherry channel. After background fluorescence subtraction and photobleaching removal, change of fluorescence was calculated using ΔF/F_0_ = (F – F0)/F0. Baseline F0 was determined from the mean fluorescence from time interval T = 2 s and T = 2.5 s. Reference ΔF/F0 was digitally filtered using lowpass and highpass filters (*lowpass* and *highpass* functions) at three ranges: [10, 30] Hz, [1, 10] Hz, and below 10 Hz. JEDI-1P-Kv ΔF/F_0_ was filtered using lowpass filter at 70 Hz. In Step 1, reference ΔF/F_0_ r_1_ was scaled and subtracted from JEDI-1P-Kv ΔF/F_0_ g_1_ and formed g_2_. In Step 2, r_2_ was scaled to and subtracted from g_2_ to form g_3_. The same regression process scaled and removed r_3_ from g_3_. The same process can be repeated until shared noises at different frequency ranges have been removed. To compare JEDI-1P-Kv signal before and after the step-by-step regression, we calculated the mean power spectral density of the signal before and after the regression step (n = 25) with fast Fourier transform (FFT) using MATLAB Spectrum Analyzer. The same power spectral analysis was used to compare the frequency distribution of JEDI-1P-Kv and reference channels.

#### Simultaneous widefield imaging and acute local field potential recording in lightly anesthetized mice

For simultaneous widefield imaging and acute local field potential (LFP) recording, mice first received the imaging window and head-post implant and were allowed at least a week to recover. On the day of recording, the animals were anesthetized with 2% isoflurane and kept on a 37°C heating pad while a small craniotomy (1 × 2 mm) was carefully drilled above the barrel cortex at the edge of the imaging window (**Fig. 5a**). After the skull was carefully lifted and the cortex was exposed, a thin layer of Dura-Gel (Cambridge Neurotech) was applied over the dura to prevent the exposed area from drying. Another small craniotomy was created above the cerebellum in order to place a stainless-steel grounding wire.

The isoflurane was reduced to 1% during the recording, and the animals were kept on the 37°C heating pad and head-fixed throughout the procedure. A 32-channel silicone probe (ASSY-37H8B, Cambridge Neurotech) was inserted into the craniotomy above barrel cortex at a 45-degree angle to 1.0 - 1.2 mm deep. The LFP data were acquired through RHD2000 USB Interface Board (Intan Technologies, LLC) at 20 kS/s, and the imaging data were acquired through MiCAM ULTIMA at 200 Hz. Each trial with simultaneous imaging and LFP recording was 20.48 s long.

#### Data analysis of widefield simultaneous imaging and acute LFP recording

To show the relationship between voltage imaging and LFP, we calculated the correlation coefficient between the two types of data (n = 3 mice). Trials with high-amplitude noise in LFP recording caused by respiration or motion were excluded from the analysis. First, we identified the channel of the neural probe that best correlated with imaging signals. LFP signals were down sampled to 200 Hz to match the imaging acquisition rate and lowpass filtered at 70 Hz. We sampled 8 ROIs from each mouse and calculated the correlation coefficient between the regressed JEDI-1P-Kv trace at each ROI and matching LFP signals from all 32 channels using MATLAB function *corrcoef*. The correlation coefficient between the same JEDI-1P-Kv signal and all electrode channels varied based on the depth of electrode channels in the cortex. For each set of matching trials with the same probe insertion depth, the modal electrode channel with maximum correlation coefficient among all 8 ROIs was identified. The modal channel number was used for the following analyses.

To understand the spatial relationship between LFP signal and cortex-wide imaging signals, we calculated correlation coefficient between the LFP and JEDI-1P-Kv signal at each pixel. Within each animal, we averaged pixel-wide correlation coefficient across all trials to form the two-dimensional heat map that shows the correlation coefficient of the entire cortex (**Fig. 5d**). To explore the relationship between LFP and imaging signal at selected ROIs, we averaged the correlation coefficient at each ROI from all animals and compared it with shuffled controls. The shuffled controls were generated by randomly mismatching LFP and imaging trials of 172 trials from all mice. A Mann-Whitney U test was used to compare the correlation coefficient at selected ROIs from matched trials with that from shuffled controls. With the same imaging and LFP dataset, magnitude-squared coherence was calculated using MATLAB function *mscohere* to show the similarity of the two types of data in the frequency domain.

#### *In vivo* imaging with sensory stimulation

We applied air-puff or light flicker stimulations to mice during widefield imaging to test JEDI-1P’s ability to follow fast frequency responses in the cortex. The animals were head-fixed during the imaging. All trials were 20.48 s long. For air-puff stimulation, the air was delivered via a metal tube placing vertically under the right-side whiskers and was controlled through a miniature solenoid valve (LHDA1233215H, Lee Company). For light flicker experiments, a white LED was placed 20 mm away from the animals’ eyes with an illuminance of 400 Lux ^86^(49). Each imaging trial started with a 5 s baseline with no stimuli and followed by stimulation at 25 (or 20 for light flicker), 30, 40, 50, and 60 Hz with 50% duty cycle that lasted 3 s at each frequency.

For experiments showing stimulation response to sensory stimuli during long-term imaging, mice received a 0.75 s air-puff stimulation at either 30 Hz or 50 Hz with 50% duty cycle. The imaging was 10.24 s per trial.

#### Statistical analysis

For each of the comparison we made in the manuscript, statistical details were specified either in the main text or along with the figure captions, including (1) the statistical test used, (2) sample sizes *n*, (3) what *n* represents, (3) the definition of the center (i.e., mean or median) and (5) the definition of the error bars. A comparison is defined to be statistically significant if the p-value was less than 0.05, unless otherwise stated.

For *in vitro* experiments, we performed two-tailed t-tests when comparing the means between two groups and ANOVA when comparing the means among more than two groups. Prior to t-test, one-way and two-way ANOVA, we first conducted F-test, Brown-Forsythe test and Spearman’s test, respectively, to test for equal variances between groups. When the variances were statistically different, the Welch’s correction was applied. Following one-way ordinary ANOVA, Welch’s ANOVA and two-way ANOVA, we conducted Tukey’s Hones Significant Difference (HSD), Dunnett’s T3 and Holm-Šídák post hoc test respectively for multiple comparison. Because normality tests have little power when the sample size is small ^87,88^(50,51), we did not perform normality check and assumed normal distribution when appropriate.

For linear regression we did for *in vitro* experiments, statistical details were specified either in the main text or along with the figure captions, including (1) how regression was performed, (2) sample sizes n, (3) what n represents, Result was reported as r-squared effective size (Pearson’s r^2^).

For *in vivo* experiments, we performed Mann-Whitney U test when comparing the experimental group and control group. Sample size n and p-value for each test were specified in the caption. Functions sigstar() and stdshade() were adapted for plotting significance stars and 95% CI, respectively ^89,90^.

## Acknowledgements

We acknowledge Dr. B. Arenkiel, J. Ortiz-Guzman, and Z. Chen at the TCH Neuroconnectivity Core for AAV packaging; this Core is supported by NIH grant P50HD103555 and the Charif Souki Fund. We thank J. M. Kirk, H. Johnson and the Optical Imaging & Vital Microscopy (OiVM) Core at Baylor College of Medicine for assistance with the confocal microscopy and the plate reader. We thank F. A. Blanco, C. A. Cronkite, and Dr. J. G. Duman from Dr. K.R. F. Tolias lab (BCM) for preparing neurons for *in vitro* characterization. We acknowledge Dr. Shella Keilholz from Emory University and Dr. Arthur Morrissette for helping with the *in vivo* imaging analyses. We thank Madison Cohen, Ellie Jiayi He, and Brune Le Chatelier at Emory University for helping with animal training.

## Authors’ contributions

DJ and FSP conceived and oversaw the project. **GEVI screening and *in vitro* characterization**: ZL developed the screening platform hardware; ZL and XL. developed the screening platform software and data analysis code; XL and YG screened GEVIs; XL characterized GEVIs *in vitro* and cloned viral vectors. XL, ZL, and FSP prepared figures and wrote the manuscript. ***In vivo* imaging**. YM conducted all *in vivo* imaging experiments and developed code for pre-processing pipeline and related analyses. ZL provided advice for the photobleaching correction. YM and DJ prepared figures and wrote the manuscript.

## Data availability statement

All source data, materials and computer code are available to all editors, reviewers and researchers upon request. The sequence of JEDI-1P is available from GenBank (accession number TBD). Plasmids used for *in vitro* characterization and packaging AAV for *in vivo* voltage imaging are available from addgene (accession numbers are pc3-puro-CAG-JEDI-1P-P2A-mCherry-CAAX, TBD; pc3-puro-CAG-JEDI-1P, TBD; pAAV-EF1a-DIO-JEDI-1P-Kv, TBD; pAAV-EF1a-DIO-mCherry, TBD; pAAV-EF1a-DIO-tdTomato, TBD. The code for *in vivo* imaging analysis is available at [Github code will be provided prior to publication].

